# Exploring the tumor micro-environment in ovarian cancer histotypes and tumor sites

**DOI:** 10.1101/2023.10.07.561344

**Authors:** Bingqing Xie, Susan Olalekan, Rebecca Back, Naa Asheley Ashitey, Heather Eckart, Anindita Basu

## Abstract

Ovarian cancer is a highly heterogeneous disease consisting of at least five different histological subtypes with varying clinical features, cells of origin, molecular composition, risk factors, and treatments. While most single-cell studies have focused on High grade serous ovarian cancer, a comprehensive landscape of the constituent cell types and their interactions within the tumor microenvironment are yet to be established in the different ovarian cancer histotypes. Further characterization of tumor progression, metastasis, and various histotypes are also needed to connect molecular signatures to pathological grading for personalized diagnosis and tailored treatment. In this study, we leveraged high-resolution single-cell RNA sequencing technology to elucidate the cellular compositions on 21 solid tumor samples collected from 12 patients with six ovarian cancer histotypes and both primary (ovaries) and metastatic (omentum, rectum) sites. The diverse collection allowed us to deconstruct the histotypes and tumor site-specific expression patterns of cells in the tumor and identify key marker genes and ligand-receptor pairs that are active in the ovarian tumor microenvironment. Our findings can be used in improving precision disease stratification and optimizing treatment options.

## Introduction

Ovarian cancer is the second most common and most malignant cancer in the female reproductive tract. According to the American Cancer Society, 90% of ovarian cancer originated from epithelial tissue and can be further divided into serous, endometrioid, clear cell, and mucinous histotypes^1^. The risk factors of epithelial ovarian cancer vary from each histotype but generally include age, weight, hormone therapy after menopause, as well as family history^2^. Previous genomic studies^3^ on ovarian cancer have investigated the effects of variations in genes that included TP53, NF1 and BRCA1. Mutations in TP53 and NF1 and dysfunction of BRCA1 are related to the pathogenesis of the serous carcinoma in ovary^4^. However, the molecular mechanism for ovarian cancer remains unclear and targeted therapy is yet to be developed. In recent years, the development of single-cell technology allows researchers to zoom in on the cell-level transcriptome of the tumor tissue and provides a better understanding of the tumor microenvironment (TME). Single-cell technology has been applied to ovarian cancer previously on malignant abdominal fluid (ascites) associated with High grade serous ovarian carcinoma (HGSOC) histotype^6^. The stress associated chemo-resistance in solid tumors from metastatic sites with HGSOC was investigated together with stroma signaling to provide insight into chemotherapy resistance^7^. A recent study used scRNA-seq on primary and untreated peritoneal metastatic site^8^ to study cancer recurrence. However, comparisons across multiple sites and histotypes are yet to be performed. We previously reported the cellular composition of metastatic ovarian tumors using single-cell RNA sequencing technology^9^. We found heterogeneity in the immune responses of different ovarian cancer patients, among immune sub-populations identified from the metastatic samples, allowing us to separate tumors into two groups based on T cell infiltration. The metastatic samples can be grouped into high and low T infiltrated types based on both immunohistochemistry (IHC) and single-cell transcriptomic profiles. We established a comprehensive collection of immune cells from the differential expression of marker genes.

In the current study, we characterized tumors from 12 ovarian cancer patients using Drop-seq, a high-throughput single-cell RNA-seq technique^5^. We broadened our focus to include primary tumor sites and other histotypes besides HGSOC which allowed us to identify cell types that are specific to sites or histotypes. We analyzed the distribution of cancer-associated cells and elucidated cell-cell communication in each histotype. We identified a cluster of cancer stem cells (CSCs) within the epithelial cells, based on their increased expression of markers *IFIT1*, *IFIT2*, *IFIT3* and *ISG15*. This cluster is present in HGSOC and MMMT histotypes. Within the stromal cells, we found multiple cancer-associated fibroblast (CAF) sub-clusters which showed high expression of *IL6*, *CCL2*, *S100A4*, *PDPN*, and *FGF7* in both primary and metastatic samples. We also verified that our previous observations on the immune cell activity in metastatic samples are still valid across a larger sample collection that includes primary tissues and multiple histotypes. In addition, we identified a cluster of IL32+ plasma B cells that were found exclusively in the primary tumor sites.

With the inclusion of additional histotypes and tumor sites in our collection, this study allows us to characterize the differences in cell compositions between sites and different levels of their T cell infiltration, build cell or gene signatures to characterize the different ovarian cancer histotypes, and further investigate the underlying molecular mechanism in the TME. We further explored cell-to-cell communication among different cell sub-clusters, using inferred ligand-receptor (LR) interactions. We note that such interactions are enriched among epithelial cells and fibroblasts and that LR interaction signatures vary across different tumor sites and histotypes.

## Results

### Establishing cell lineages, TCGA subtypes and cell-cycle states across samples

To study cell composition of ovarian cancer, tumor tissues resected from 12 ovarian cancer patients undergoing debulking surgery in the ovaries, omentum and rectum were analyzed in this study (**Fig. 1A**, **Table 1A**). Briefly, the cohort consisted of seven white, two Asian, two Black women and one woman of unknown racial origin and ranged between 39-77 yr in age (mean ∼62 yr). Most patient tumors were stage IIIB or above according to staging by a pathologist. Solid tumor samples of different histotypes were collected from primary (ovaries) and metastatic (omentum, rectum) sites (**Table 1B**) which enabled us to investigate histotype- and site-specific signatures at single cell level. Tumor samples were obtained fresh from surgery and processed using Drop-seq^5^ within 24 hr or fixed in formalin for immunohistochemistry (IHC). Immune (CD45+) and cancer cells enriched from a subset of samples were also profiled by Drop-seq to obtain better representation of immune cells in our single cell data.

**Figure 1:**
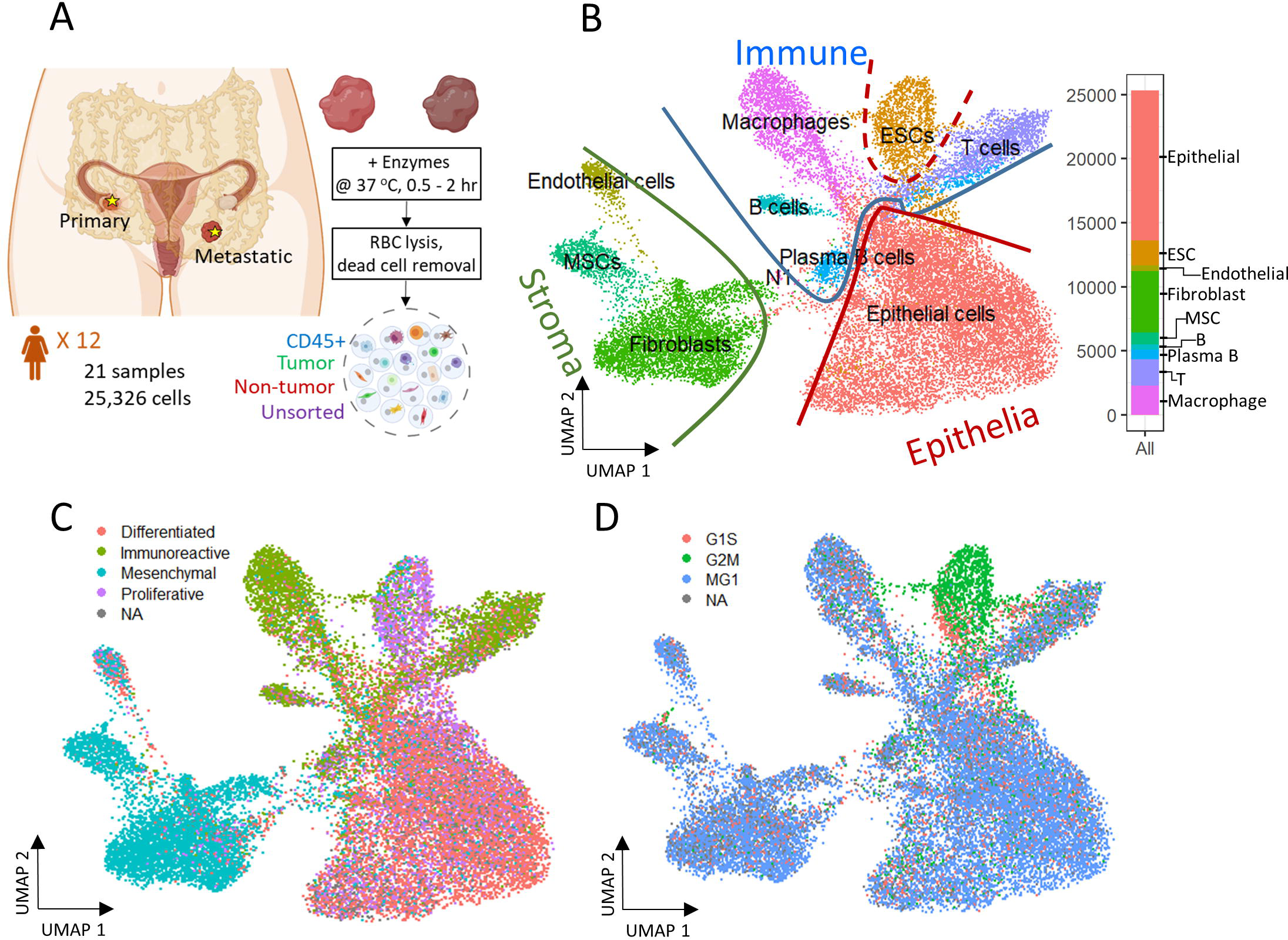
Experiment design and 2D reduced representation of all cells included in the study, annotated by major cell lineage, predicted cancer subtype and cell cycle phase. A) Profiling ovarian cancer tumor samples of different using droplet single cell RNA-seq. B) All cell types projected on UMAP divided by Epithelia, Immune and Stroma subpopulations. C) Predicted cancer subtype projected on UMAP. D) Cell cycle assignment projected on UMAP.

**Table 1:**
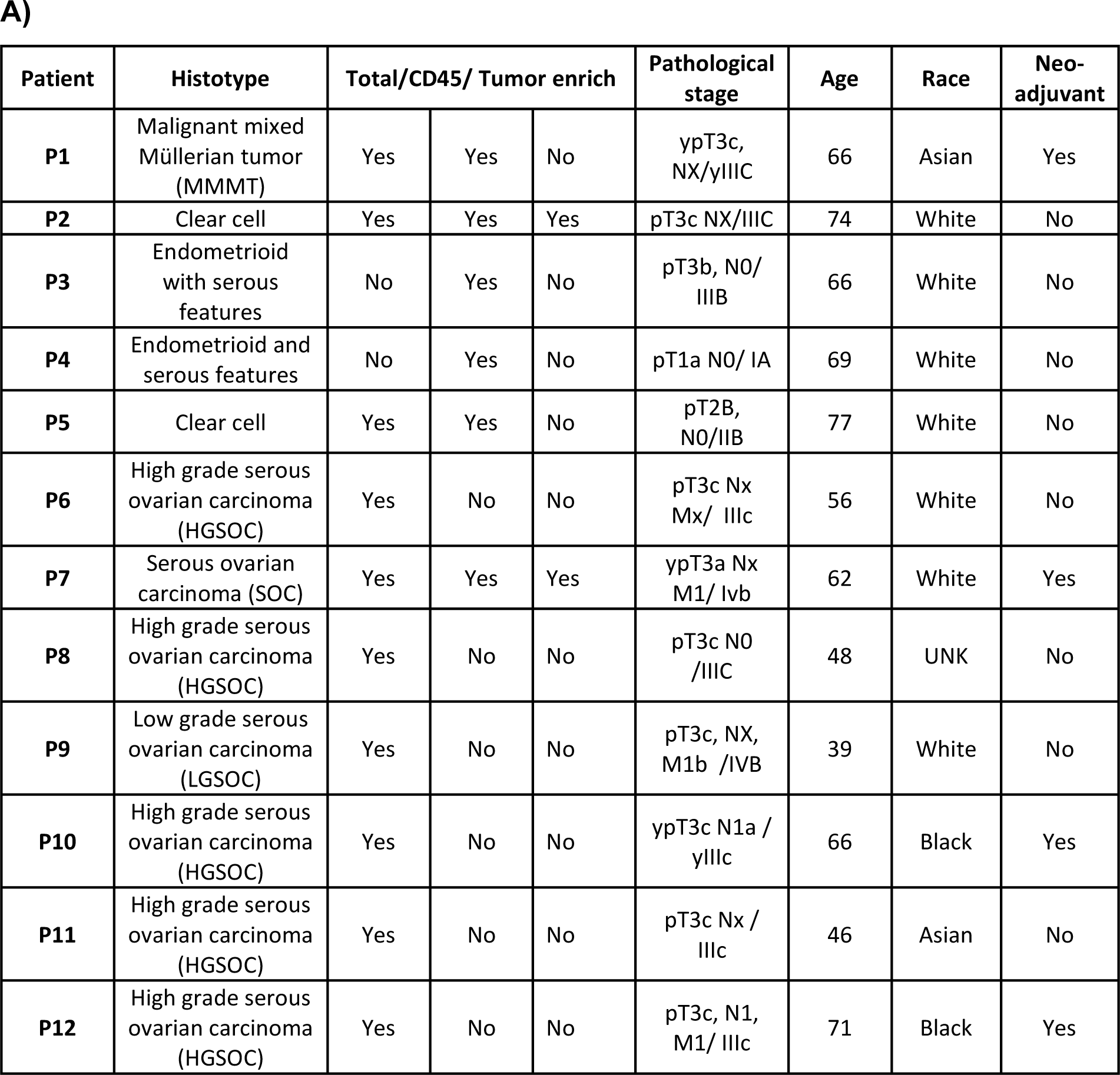
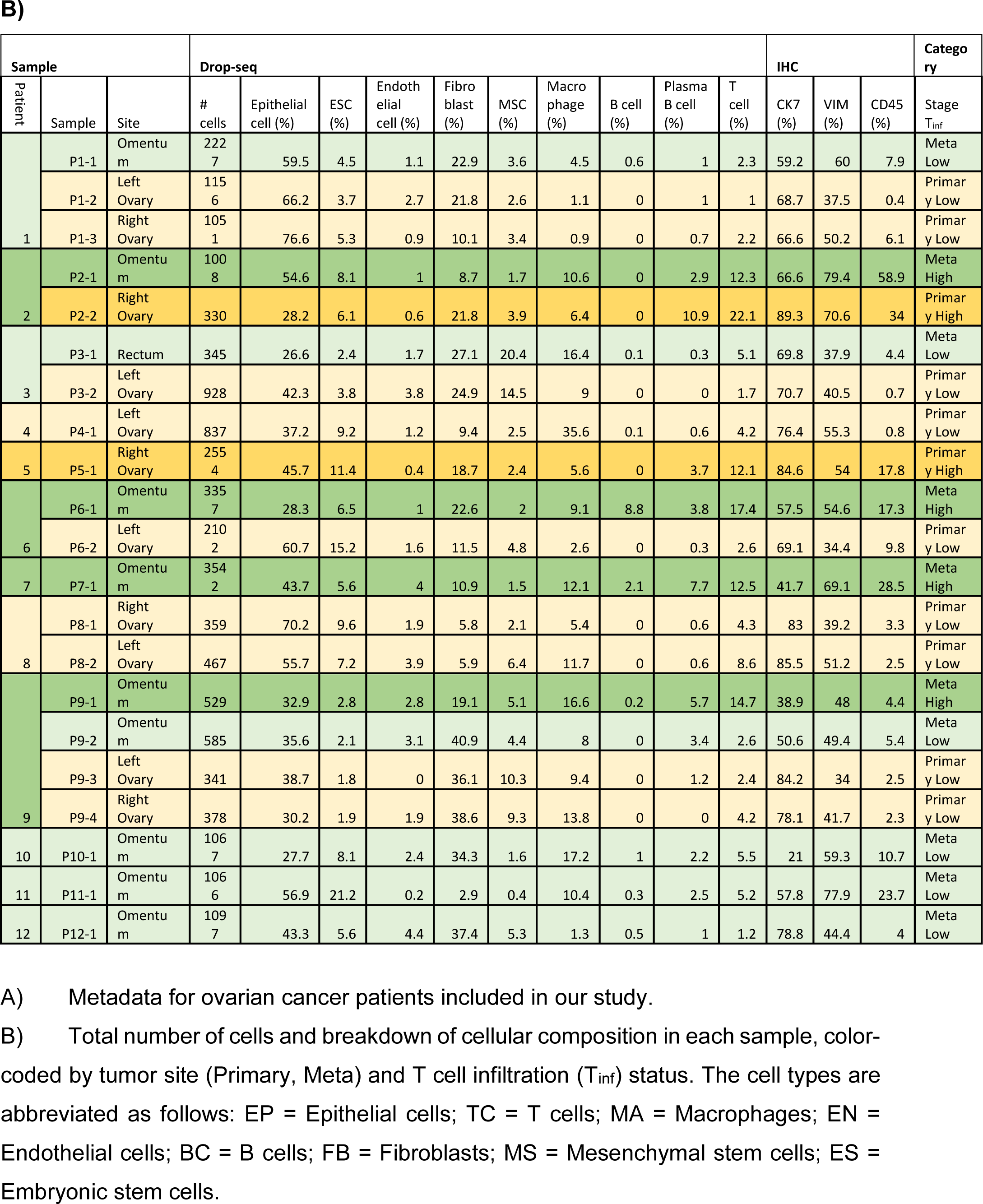

A total of 26 gene expression matrices were generated from Drop-seq experiments on 21 ovarian cancer tumors from 12 patients. We identified a total of 38,811 genetic features across 25,326 cells from tumors resected from multiple tissue sites in this study. The filtered gene expression matrices were integrated using the anchor-based alignment. Unsupervised clustering analysis yielded 11 distinct clusters of cells. The resulting clusters were annotated using Template-based Automated Cell type Assignment (sc-TACA; **Methods**), yielding ten major cell types including epithelial, endothelial, mesenchymal stem (MSC), embryonic stem (ESC), fibroblast, macrophage, T, B and plasma B cells and a small cluster of 37 cells marked as N1 that shared markers with astrocytes which we saw in our previous study^9^ (**Fig. 1B**). Percentages of each cell type comprising each tumor sample are shown in **Fig. S1A** and **Table 1B**. Due to the small number of N1 cells in any given sample (< 0.1%), we excluded them in further analysis. For simplicity, the cell-types were classified into three compartments: *epithelia,* containing epithelial cells and ESCs, *stroma* containing endothelial cells, MSCs and fibroblasts, and *immune*, containing macrophages, B and plasma B cells, and T cells (**Fig. 1B**).

Next, we explored the expression of the genes associated with the four molecular subtypes of ovarian cancer-differentiated, immunoreactive, mesenchymal and proliferative- identified by TCGA^3^ in our dataset. We were able to assign one of the four molecular subtypes with the highest TCGA module score to 93.7% of cells; cells with a negative module score were marked as not assigned (NA)^9^. When each cell on the UMAP was marked with the molecular subtype assigned to it (**Fig. 1C**), we noted that the major cell types and the cellular compartments they belong to (**Fig. 1B**) match the predominant molecular subtype of ovarian cancer identified by TCGA. The epithelial cells were distributed through all four cancer subtypes and comprised 80% of the cells predicted as differentiated subtype. 73% of cells from the predicted immunoreactive subtype were immune cells (B cells, T cells, and macrophages). The mesenchymal subtype, associated with worst survival^10^, consisted of the least epithelial cells and contained the highest percentage (82%) of stroma cells, including MSCs, fibroblasts, and endothelial cells. The proliferative subtype contained 56% cells from the epithelial cell category; 26.2% cells from the ESC (about half of the total ESC population) that also showed unique stem cell features described later, were of the proliferative subtype. Sample-specific composition of TCGA subtypes is shown in **Fig. S1B**.

To study the cell cycling effects under the TME, Cell cycle analysis was performed on the combined dataset to assign a cell-cycle module score to each cell for the G1S, G2M and MG1 phases. Cells that could not be assigned to one of these phases were marked as ‘NA’. We noted that the cell cycling patterns were roughly similar for all cell types (**Fig. 1D**), with the exception of ESCs. A large fraction of cells across all cell types were assigned to the MG1 phase (64.3%; **Fig. S1C**), as seen previously^11^. In contrast, most ESCs (>70%) were assigned to the G2M phase where they likely stalled during the cell cycle^12^.

### Immune cells and their expression in ovarian cancer samples

We identified 5,453 cells as immune cells that could be further split into B cells, plasma B cells, T cells, and macrophages (**Fig. S2A**). We also found a few dendritic cells and common myeloid progenitor cells (52 and 30, respectively) that co-clustered with macrophages and were removed for downstream analysis due to the low cell counts. When identifying the subclusters within each cell type, we denote them as ‘EP’ for epithelial cells, ‘TC’ for T cells, ‘BC’ for B cells, ‘MA’ for macrophages, ‘ES’ for ESCs, ‘FB’ for fibroblasts, ‘MS’ for MSCs, and ‘EN’ for endothelial cells. We used a single digit starting from 0 to index the sub-clusters for each cell type, e.g., EP0 denotes cluster 0 of epithelial cells.

To determine if there were any cells unique to the different tumor sites, we cross-referenced 5,371 immune cells with our previous study^9^ of metastatic ovarian tumors. We identified five subclusters (**Fig. 2A**), consisting of three clusters of CD4+ T helper (T_h_) cells (TC0, TC3, TC4), and two clusters of CD8+ resident memory T (T_rm_) cells (TC1 and TC2). Among these clusters, one subcluster containing GNLY+CD8+T_rm_ cells (TC2) that was not observed previously^9^, derived from metastatic samples, P5-1, P6-1, P7-1. Three PRDM1+JCHAIN+ plasma B clusters, BC0, BC2, and BC3 and one naïve B cluster, BC1 were observed (**Fig. 2B, Fig. S2B**). The BC2 subcluster that consisted of CD38-SDC1-S100A4+IL32+GAPDH+ plasma B cells, has not been identified previously^9^. The marker *IL32* was a proliferation marker for malignant plasma cells in myeloma^13^. Intriguingly, we find BC1 to be almost absent in the primary tumor site (ovary). Four macrophage subclusters (**Fig. 2C, Fig. S2B**) were annotated, including a CD14+MSR1+CD163-cluster (MA1) that were mainly found in samples collected from the primary tumor site (ovary) and thus not seen in our previous study^9^.

**Figure 2:**
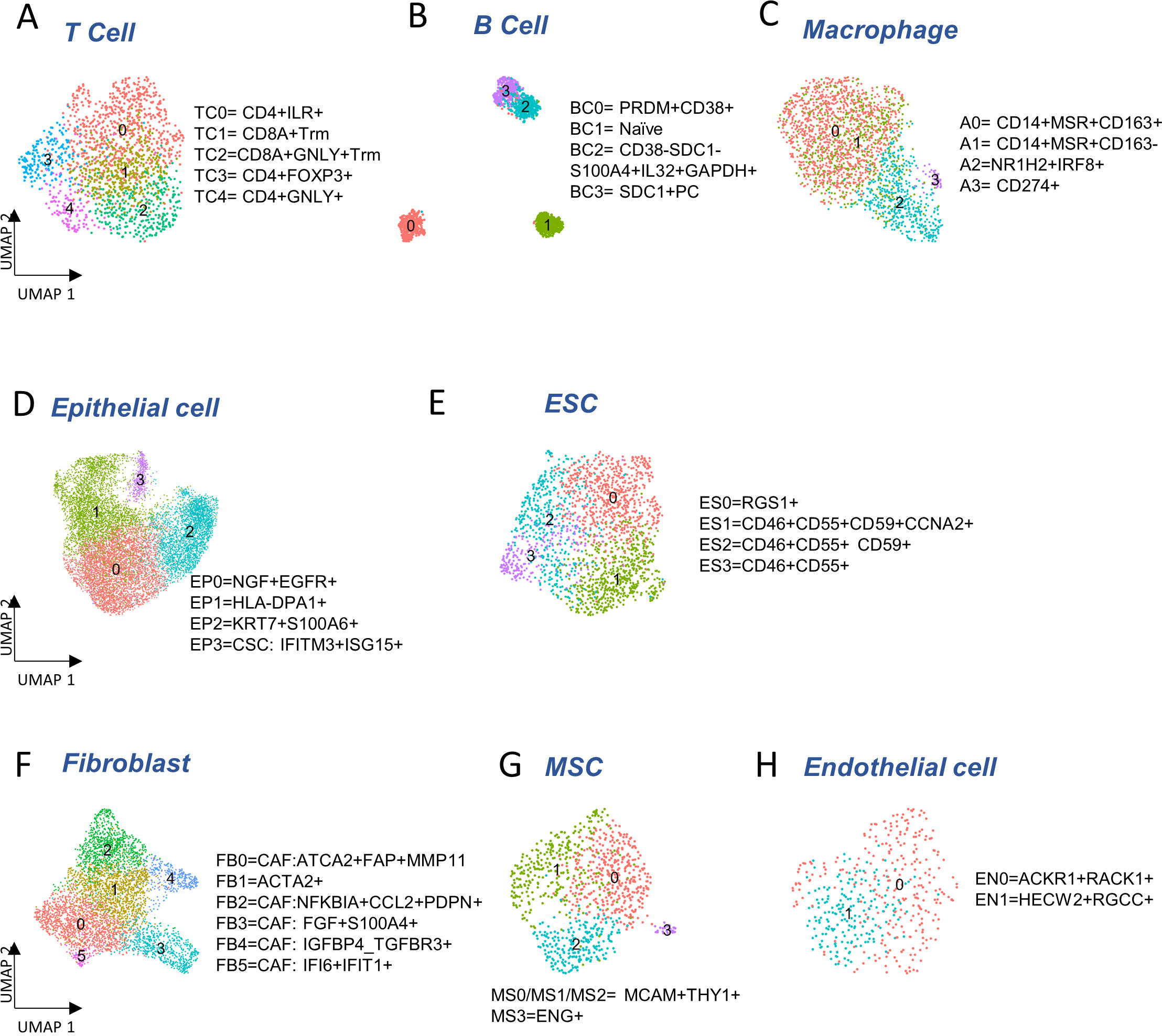
Cellular sub-types in the Immune, Epithelia and Stroma. A-C) Subclusters of major immune cell types: T cells, B cells and macrophages, respectively. D-E) Subcluster annotation for epithelial and embryonic stem cells (ESC), respectively. F-H) Subcluster annotation for major cell-types in the stroma: fibroblast, mesenchymal stem cells and endothelial cells, respectively.

### Epithelial cells and their expression in ovarian cancer samples

We detected 11,716 epithelial cells comprising the *epithelia*, as the most abundant cell type in our integrated and batch corrected dataset. Hierarchical clustering of these cells (resolution = 0.3) detected four epithelial sub-clusters, EP0-3 (**Fig. 2D**). Dot plots for some top differentially expressed markers, *EPCAM, S100A1, KIAA1217, MAML2, MECOM, IFIT2/3, and LIPA* are shown in **Fig. S2C**. The EP0 sub-cluster comprised 38% of all epithelial cells and showed a distinctive signature of cytokeratin genes, *KRT19* (logFC = 0.96), *KRT18* (logFC = 0.747) and *KRT7* (logFC = 0.633) (**Table S1A**). A recent study on the origin of ovarian cancer^14^ connected fallopian tube epithelial cell subtypes to intra-tumor heterogeneity in serous ovarian cancer (SOC), and used *KRT7* as a marker for secretory epithelial (SE) cells in the fallopian tube as the cell-of-origin for SOC. Other genes found upregulated in EP0 (**Table S1A**) were *S100A6* (logFC = 0.81) and *S100A11* (logFC = 0.66) from the S100 calcium-binding protein family. The S100 protein family interacts with cytoskeletal proteins^15^ and may promote metastasis and stimulate angiogenesis. Specifically, *S100A11* gene^16^ acts as a tumor promoter by regulating MMP activity and the epithelial-mesenchymal transition (EMT) process. Another top expressing marker gene, *LGALS3* (logFC = 0.8) is associated with cell migration, proliferation, adhesion, cell-cell interaction in tumor cells, and implicated in tumor progression and chemo-resistance of epithelial ovarian cancer^17^. EP1 cluster exhibited significant upregulation of genes belonging to the MHC class II protein family, *HLA-DPA1*, *HLA-DRA*, *HLA-DPB1*, and *HLA-DRB1* (logFC > 0.96), was associated with the KRT17 sub-cluster of secretory epithelial cells in the fallopian tube epithelia^18^ as well as high expression of ribosomal proteins such as *RPLP1* (logFC = 0.84) and *RPS6* (logFC = 0.81). EP2 subcluster was enriched for chromatin pathways, growth factor signaling pathways, such as platelet-derived growth factor (PDGF), nerve growth factor (NGF), epidermal growth factor receptor (EGFR), platelet-derived growth factor receptor Beta (PDGFRB) and angiopoietin like protein 8 (ANGPTL8) regulatory pathways. Protein families with ankyrin-repeat proteins and zinc finger proteins associated with cancer progression^19,20^ were upregulated in EP2. EP3 showed a unique signature of interferon-stimulated genes *IFIT1-3* (logFC > 2.5), *IFITM1-3* (logFC > 0.6), and *ISG15* (logFC = 2.5), previously characterized as markers of cancer stem cells (CSC)^21^. Detailed marker information is provided in **Table S1A**.

We also detected 1,925 embryonic stem cells (ESC) in our combined dataset that showed moderate expression of ESC markers, *STAT3* and *CTNNB1* (**Fig. S2D**). Further clustering of the ESCs yielded 4 sub-clusters (**Fig. 2E**): ES0 exhibited markers of the immunoreactive molecular subtype, such as *RGS1*^22^ (logFC = 1.91), *CD3E*^23^ (logFC = 0.55) and *CD3G*^23^ (logFC = 0.82); also see **Table S1C**. We found cancer stem cell gene, *CD24*^24^ and therapy resistant genes, *CD46* and *CD55*^25^ expressed in ES1, ES2, and ES3 and cancer stem cell marker, *CD59*^26^ in ES1 and ES2 (**Fig. S2D**). Analysis of cell cycle activity (**Fig. 1D**) assigned 73% of the ESCs to the G2M and 22.5% to the G1S phases. The elongated G2M phase has been previously associated with cancer cell proliferation, mutation of *TP53* and *KARS*, T cell infiltration, and cancer metastasis^27–29^. Moreover, the expression of CDKN1A and senescence gene FN1 with the lack of expression of PCNA can trigger the G2 arrest or the stress-induced premature senescence (SIPS) found in a previous cancer study^30^ (**Fig. S2D**). Meanwhile the shortened G1 phase regulated by *TP53* can lead to DNA damage, and subsequently affect the S phase with malfunctioned G1/S checkpoint^31^. Low number of cells in MG1 (< 5%) may indicate the ESCs to be post-mitotic. Specifically, genes expressed in ES1 are enriched for cell cycle functions and G2/M transition (**Table S1D**). Increased gene expression required for G2/M transition and indicative expression for DNA damage response, such as *CCNA2, CCNB1, CCNB2, CDK1, CKAP5, DCTN3* and *TUBB4B* can support the malfunction of P53^32^ (**Fig. S2D**).

### Stromal cells and their expression in ovarian cancer samples

The second largest cellular compartment, *stroma*, contained three major subsets: fibroblasts, MSC and endothelial cells. Hierarchical clustering of 4,772 fibroblasts (resolution = 0.5) yielded six sub-clusters (**Fig. 2F**) containing markers for cancer-associated fibroblasts (CAF). The CAF-like clusters involved multiple molecular mechanisms associated with tumor progression, angiogenesis via vascular endothelial growth factor A (*VEGFA*) production, and coordination of immune function through chemokine and cytokine^33^ production. FB0 and FB1 showed comparatively high expression of myofibroblast markers, *ACTA2* (logFC = 1.04) and *MYL9*^34^ (logFC = 0.74). CAF associated markers, *MMP11*, *MMP2*, *FAP*, *THY1*, *IFI6*, *IFI27*^35^ (logFC > 0.33) were highly expressed in FB0 (**Fig. S2E, Table S2A**). In contrast, we did not find any CAF-related expression in FB1. FB2 showed upregulation of NF-kappa B signaling pathway genes, *NFKBIA*, *NFKB1* and *NFKBIZ* (logFC > 0.42), VEGFA-VEGFR2 signaling pathway gene, *VEGFA* (logFC = 0.33), chemokine receptor genes, *IL6* (logFC = 1.7) and *CCL2* (logFC = 1.72), transmembrane glycoprotein genes, *PDPN*^36^ (logFC = 0.1), and genes associated with cancer metastasis, *IER3*^37^ (logFC = 0.633), *SGK1*^38^ (logFC = 1), and *SERPINE2*^39^ (logFC = 0.5). Genes overexpressed in FB3 subcluster were enriched for angiogenesis, integrin signaling and related to extracellular matrix remodeling, including *FGF7* (logFC = 1.18) and *S100A4* (logFC = 0.89). The FB4 subcluster exhibited elevated expressions of growth factor binding genes, *IGFBP4* (logFC = 0.83), *TGFBR3* (logFC = 0.82) and top markers *APOLD1*, *MCAM*, and *PLXND1* for angiogenesis and blood vessel development (**Tables S2A, B**). FB5 showed upregulated genes highly enriched in immune crosstalk and cytokine/interferon signaling pathways. Particularly, interferon inducible genes, such as *IFI6*, *IFI27*, *IFI44*, *IFI44L*, *IFIH1*, *IFIT1-3* (logFC > 1.53) were highly expressed in FB5 subcluster that might be due to the inflammatory crosstalk in the TME^40^ (**Table S2A, B**). Taken together, all fibroblast subclusters exhibited CAF features, with the exception of FB1.

The progenitors of stroma sub-population, 951 mesenchymal stem cells (MSC) were detected in our data that could be clustered into 4 sub-clusters (**Fig. 2G**). The majority of the MSCs (MS0-3) expressed MSC markers, *MCAM* and *THY1* (**Fig. 2G, Fig. S2F**). A small subset of MSCs (MS3) also expressed *ENG* that was not seen in the other clusters (**Fig. S2F**).

Finally, a distinct population of 472 endothelial cells was found in the stromal compartment. Two sub-clusters, EN0 and EN1, both expressing endothelial markers, *ENG*, *S100A6*, and *CD34*^41–43^ were found (**Fig. 2H**). EN0 showed higher expression of *ACKR1* (logFC = 1.22) which is associated with ligand transcytosis^44^ and serves as a non-specific and promiscuous receptor for several inflammatory chemokines when expressing in endothelial cells^41,45,46^, carcinoma-associated genes, *RACK1*^47^ (logFC = 0.7) and *CD74*^48^ (logFC = 0.84) (**Fig. S2G**, **Table S3A**). Genes upregulated in EN1 are related to angiogenesis and blood vessel morphogenesis in tumor metastasis (**Table S3B**).

### Cellular composition by ovarian cancer histotypes and tumor sites

We conducted further analysis on our tumor samples to examine cell types described above (**Fig. 1B, Table S4A**), cancer histotypes (**Fig. 3B**, **Table 1A**) and T cell infiltration into tumors (**Fig. 3A, Fig. S2A**, **Table 1B**). Based on pathology grading, the samples in this study belong to six ovarian cancer histotypes: serous ovarian carcinoma (SOC), high grade serous ovarian carcinoma (HGSOC), low grade serous ovarian carcinoma (LGSOC), clear cell, endometrioid with serous features, and malignant mixed Müllerian tumors (MMMT). **Fig. 3B** shows the heatmap of cell type compositions combined across all samples, grouped by cancer histotypes. We noted the highest fraction of epithelial cells in MMMT and the highest fraction of MSCs in endometrioid samples. Expression of previously established immunohistochemical markers^49,50^ *WT1*, *NAPSA*, and *PGR* for histotype classification were checked on EP and ES cell lineages. We confirmed higher expression of *WT1* in HGSOC and *NAPSA* in Clear Cell histotypes, compared to other the remaining histotypes, and the presence of *PGR* expression in endometrioid with serous features (**Fig. 3C**). Additional markers^50–52^ *VIM*, *CDKN2A,* and *ARID1A* were applied to distinguish other histotypes. The EMT repressors, zinc finger E-Box binding homeobox 1 and 2, (*ZEB2, ZEB1*)^51,52^ related to endometrial carcinosarcoma, a mix of (epithelial) carcinoma and (mesenchymal) sarcoma, were used to distinguish endometrioid and MMMT histotypes. In epithelial and ES cells, we observed *WT1+CDKN2A*+ for HGSOC, and *WT1+CDKN2A-VIM+* for LGSOC (**Fig. 3C**). Among the other histotypes, we find SOC to be *WT1+CDKN2A+VIM+*, clear cell to be *WT1-NAPSA+*, endometrioid to be *WT1-NAPSA-PGR+ZEB2+ARID1A+* and MMMT to be *WT1+VIM+ZEB1+* in our limited sample space (**Fig. 3C**).

**Figure 3:**
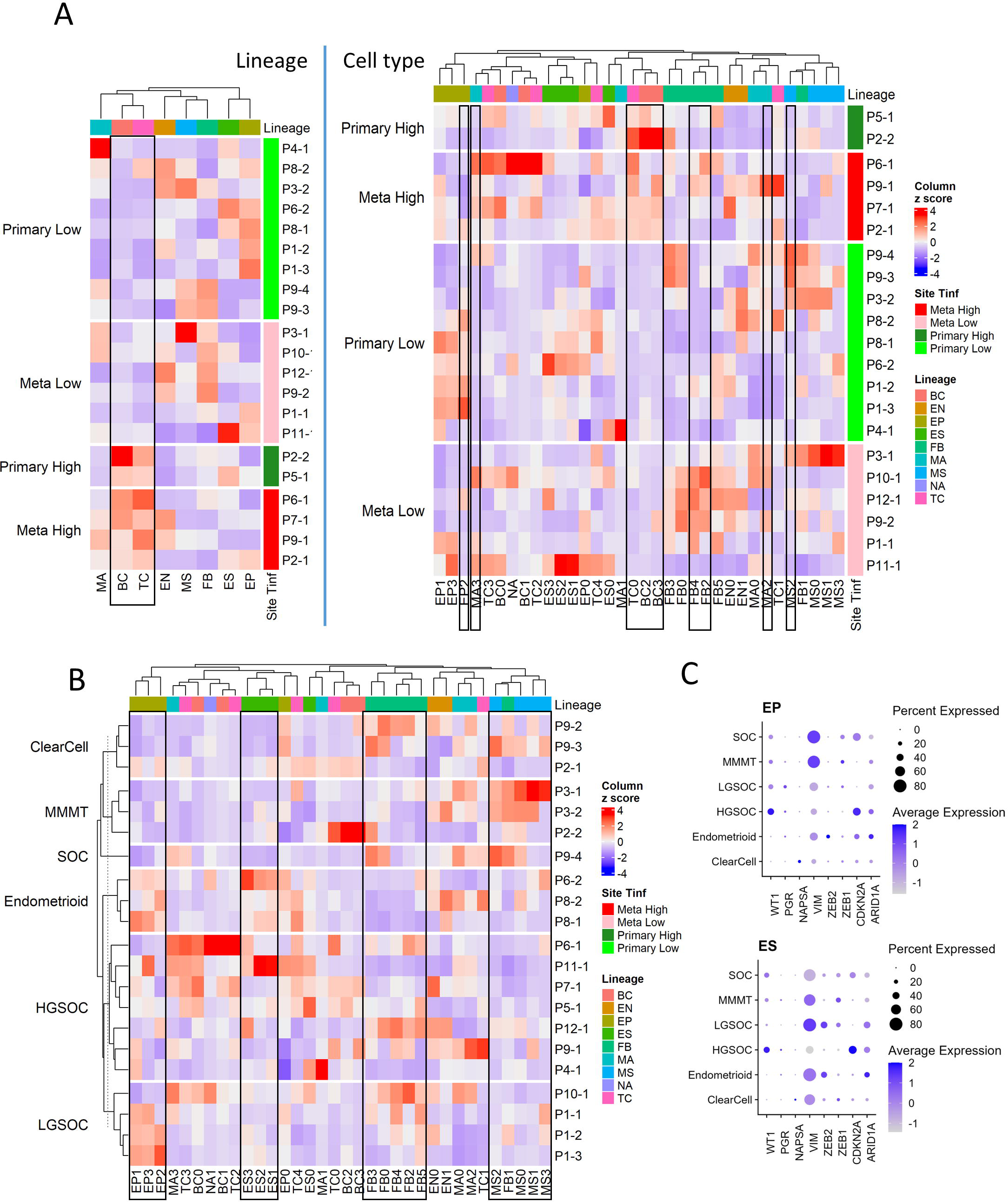
Cell composition by tumor site, T cell infiltration, and histotypes; fraction of immune, stromal and epithelial cells are explored using immunohistochemistry. A) Heatmap of major cell type composition (left) and sub cell type (right) for all patient samples. The column z-scores are calculated from cellular compositional percentages within each sample; the rows are split by site and T cell infiltration status. B) Heatmap of cell type subclusters composition percentage for all patient samples. The values are column z-scores normalizing the percentage and the rows are split by histotypes. C) Dot plot of histotype markers expression on EP and ES cells, The expression in the dot plot is the averaged scaled log normalized TP10k value.

Tumors from the ovaries were considered primary tumor sites while the tumors from the omentum and rectum were categorized as metastatic (**Table 1B**). We categorized each sample based on the level of T cell infiltration (T_Inf_)^9^, and whether its tumor site was primary or metastatic, thus grouping our samples into 4 categories: Metastatic High (T_Inf_), Metastatic Low, Primary High, and Primary Low. We found no significant differences in the composition of major cell lineages between primary and metastatic sites. However, at the sub-cellular level, the ratios of FB4, FB2, MA3, and MA2 were higher in metastatic sites, while EP2 showed an opposite pattern (**Table S4B, C**). T-tests performed on the ratios of different cell lineages showed significantly higher fractions of TC and BC in high T_Inf_ group and MS was lower in T_Inf_ (**Table S4B**). Zooming in, the TC0 and BC3 appear to drive these differences, while higher MS2 correlated with low T cell infiltration (**Fig. 3A, Table S4C**). **Fig. S2A, bottom** shows the percentages of each immune cell type in these four categories. We found FB sub-clusters, FB0, FB2, FB4 and FB5 expressing CAF markers to be enriched in samples classified as Metastatic Low (**Fig. S3B**), along with CSCs (EP3 sub-cluster, **Fig. S3B**).

Overall, the cell sub-type fractions were correlated within main cell types. For example, all EP, ES and CAF sub types-FB0, FB2, FB3, FB4 and FB5 clustered together (**Fig 3B**, black rectangles). Interestingly, the only non-CAF subcluster of fibroblast, FB1 clustered with MSC (**Fig. 3B**, black rectangle).

Due to the limited number of samples available for all histotypes, we were unable to calculate statistical significance on the cell cluster compositions. Nevertheless, several intriguing observations merit attention. The EP0 cluster was observed in all histotypes (**Table S4A)**. The fractions of EP1, EP2 and EP3 cells were higher in MMMT compared to the other histotypes, while the fraction of cells in ES1, ES2 and ES3 were higher in HGSOC (**Fig. S3A**). For clear cell histotype, the percentage of cells in TC0, BC2 and BC3 were higher. The endometrioid histotype showed a high fraction of MSCs, MA1 and FB1. The percentage of certain macrophage and T cell subtypes in LGSOC (MA0, MA2 and TC1) was higher than in other histotypes. The SOC sample of undetermined pathology grade appeared more similar to HGSOC from the primary site than LGSOC in terms of cell type composition (**Fig. S3A**). In the SOC sample, the human leukocyte antigen (HLA) genes have higher average expression compared to HGSOC samples. BC2 derived from tumors with high T cell infiltration and were identified primarily in clear cell and SOC histotypes (**Fig. S3A, B**).

Immunohistochemical staining of vimentin (VIM), CD45, and cytokeratin-7 (CK7) were also performed on tumor tissues from metastatic (**Fig. S3C**) and primary (**Fig. S3D**) tumors belonging to different ovarian cancer histotypes to investigate the fractions of the major cell lineages in these tumors. We correlated the percentage of each cellular subsets in our combined dataset from 18 samples to the IHC results; three samples from patients P3 and P4 that were enriched for CD45+ cells alone for Drop-seq were excluded from this analysis. The percentage that stained for CD45 was well correlated (Pearson correlation = 0.51 and a significant 0.03 P-value estimation) with the *immune* population (macrophage, T, B and plasma B cells). The correlations between area staining for CD45 (IHC) and percentage of T cells, plasma B cells and macrophages are 0.59, 0.53 and 0.17 (not significant), respectively. The CK7 percentage was positively correlated (Pearson’s correlation of 0.28) with the *epithelia* (epithelial cells and ESC), however not significantly (P-value = 0.24). Out of the 18 samples, only 3 samples had more than a 30% difference in percentages between the stained CK7 and annotated epithelial sub-population. The *stroma* population was estimated using the union of fibroblasts, MSC, and endothelial cells in Drop-seq data. The Pearson product-moment correlation with the percentage of cells that stained for vimentin was negative (−0.44) with a non-significant P-value and may be caused by the epithelial cells undergoing EMT (we observed a consistently smaller percentage of the stromal subpopulation compared to the VIM-stained percentage).

As seen previously^9^, we noted significant differences in the abundance of T cells between samples reported by Drop-seq. The T cell percentages in Drop-seq data showed the highest correlation with CD45 staining in IHC. Due to the correlation of T cells in tumors and cancer outcome^9^, we categorized a sample as having high T cell infiltration (T_inf_) if the percentage of T cells was greater than 10%, and low T_inf_ if less than 10% in the sample from Drop-seq data (**Fig. S2A**, **Table 1B**).

### Inferring cellular interactions in the tumor microenvironment using ligand-receptor analysis

To understand the patient-specific TME, we predicted the ligand-receptor interactions among the cell sub-clusters, using *CellPhoneDB*^53^ and additional cancer-specific ligand-receptor (LR) pairs that were curated from previous studies (see **Methods**). We found that FB, EP, ES and MS cells were highly activated for ligand-receptor (LR) interactions (**Fig. 4A**). The higher abundance of FB and EP cells in the TME and high numbers of putative LR interactions identified within and between EP, FB and immune cells in our data allowed us to further dissect histotype- or site-specific LR repertoires. Accordingly, we selected the following lineage pairs: epithelial-to-fibroblast, immune-to-epithelial, and immune-to-fibroblast. Clusters with less than 50 cells were excluded from the downstream LR analysis.

**Figure 4:**
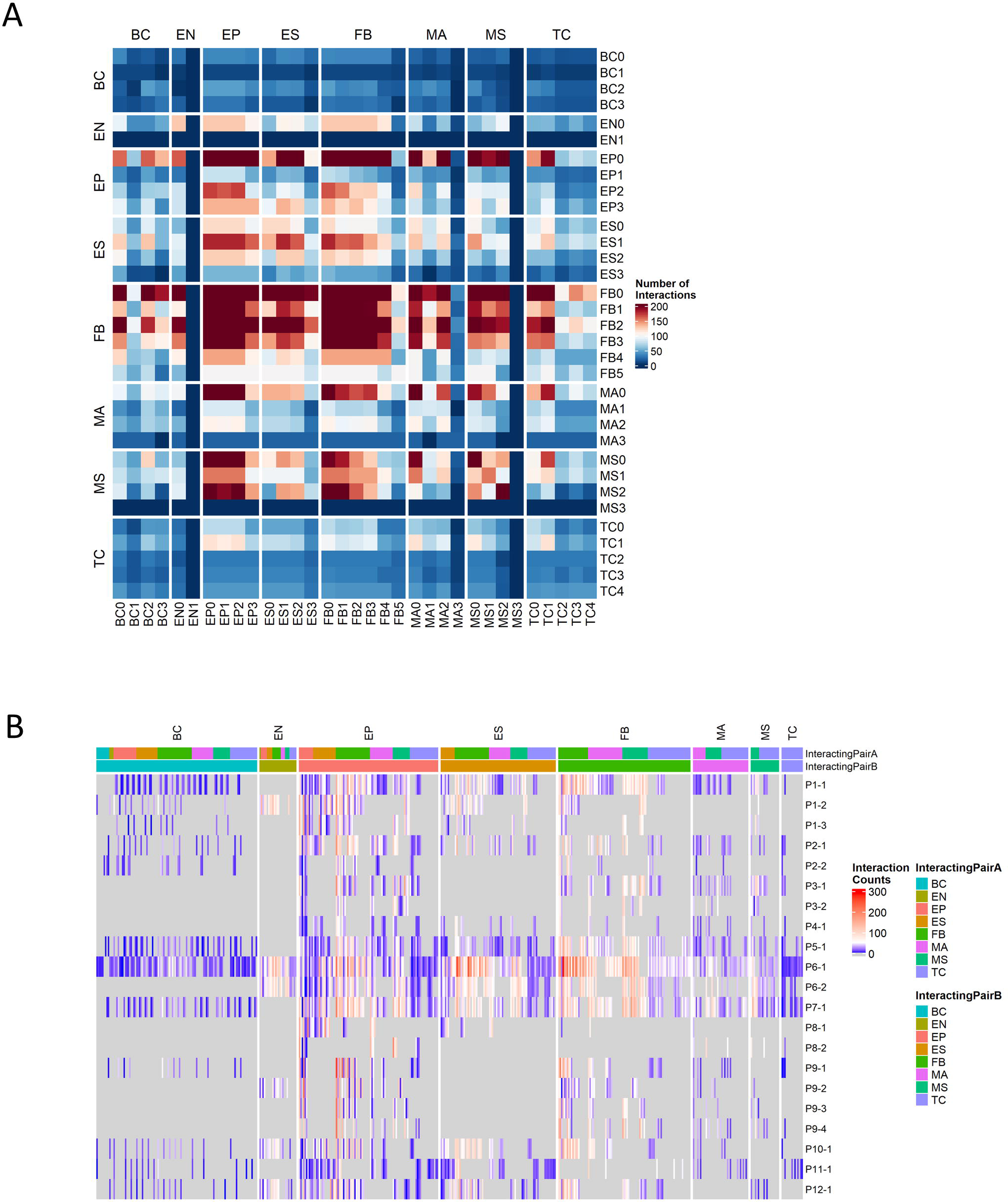
Ligand-receptor (LR) interactions predicted by CellPhoneDB using a customized cancer database. A) Total number of interactions between all cell subtypes. B) Counts of significant Ligand-receptor pairs for all cell type subclusters stratified by sample, the columns are grouped by the cell lineage of the first interactor.

We first examined the resulting cancer-specific LR interactions in epithelial-to-fibroblasts. To identify LR interactions common to each histotype, we integrated interactions from all samples grouped by their histotypes. Histotype-specific LR signatures across epithelial-to-fibroblasts were identified (**Fig. S4A**). HGSOC displayed higher interactions of receptors *ITGB1* in epithelial cells (to *COL1A2*, *MDK*, and *VEGFA* in fibroblasts), as well as *FGFR1* in epithelial cells (to *FGF12* and *FGF18* in fibroblasts). LGSOC histotype had higher LR signatures for *ITGA5*_*ADAM17*, *MET_SEMA5A*, *LAMB1*_*ITGA2*, *LAMC1*_*ITGB4*, and VEGFA_NRP2. Receptor *FGFR1* was also highly expressed in epithelial cells in LGSOC, though the ligand it enriched for was *FGF9*. Clear cell histotype has unique signatures of *CCL2_CCR3*, *SILT2_SDC1*, *BMP2_BMPR2*, and *FBN1_ITGA5*. The endometrioid histotype displayed receptor FGFR3 in fibroblast and ligands *HSP90AA1* and *FGF12* in epithelial cells. MMMT histotype showed ligand *IGF2* in epithelial cells and receptors *INSR*, *IGF1R*, and *IGF2R* in fibroblasts. Histotype with SOC features (patient P7 only) shared some LR signatures with HGSOC and LGSOC while having distinct combinations of *IGF1_IGF1R*, *SLIT2_ROBO1*, and *EFNA5_EPHB6* (**Fig S4A**).

For cancer associated LR interactions from the immune-to-epithelial (**Fig S4B**), the HGSOC patients had higher *ITGA4*_*MDK* and *BTLA* (to *VTCN1* and *TNFRSF14*). LGSOC was enriched for *THBS1* (interacting with *CD47*, *ITGA3*, *ITGA6*), and *ITGB1* (interacting with *LAMC2*, *ADAM17*, and *TGFBR2*). The SOC histotype predicted *CD44* binding to *VIM* and *FN1.* Clear cell histotype had signatures of *CCL5*_*CCR3*, *ITGA4*_*VCAN*, *CD44*_*SPP1*, *KLRD_HLA-E* and *IL2RB*_*IL15.* Endometrioid histotype had distinct signatures, such as *C1QB*_*LRP1,* and *TNF*_*LTBR*. The LR pairs for MMMT histotypes were *MMP9_LRP1, LRP1* (to *PSAP*, *SERPING1* and *A2M*), *COL2A1_DDR1* and *ITGB1_COL2A* (**Fig S4B**).

The cancer-associated LR interactions in immune-to-fibroblast subset (**Fig. S4C**) identified high number of *CD44* and *VIM* receptors interacting with *COL1A1* and *ITGB1_COL1A2* for the HGSOC histotype. LGSOC has higher *AREG_EGFR*, *INSR_NAMPT*, *EREG* and *HBEGF* to *EGFR*, *TFGB1_TFGBR3* interactions, and shared *ITGA4_THBS1* interaction signatures with HGSOC. SOC shared *ITGB1* (to *THBS2* and *LAMB1)* interactions with HGSOC. Signatures of *ITGA6* interactions with *THBS2* and *LAMB1* were higher in SOC histotype alone. *DDR2_COL1A1*, *IGF2R_IGF, VEGFB_NRP1* and *PDGFA*_*PDFGRA* were exclusively present in the MMMT histotype. Endometrioid histotype also had unique signatures, such as *CD44_LAMC3, PTPRC_CD22* and *FN1* (with *ITGA8* and *ITGA9*). The *KLRD1_HLA-E* and *ITGB7_VCAM1* were found in the clear cell histotype (**Fig S4C**).

For samples with abundant immune subpopulations, it is feasible for us to break down the immune cells into those compartments with sufficient number of cells captured. The original *CellPhoneDB* database was used to capture commonly occurring LRs that may not be specific to cancer in immune cell subpopulations. We ranked samples by the number of LR interactions (**Fig. 4B**) and selected four samples with high LR interactions for comparison: P1-1 (metastatic, low T_Inf_), P6-1, P7-1 (metastatic, high T_Inf_), and P5-1 (primary, high T_Inf_). In particular, we examined LR interactions of T cells with fibroblasts (**Fig. S5A**) and ESCs (**Fig. S5B**). We observed common signals for *TIGIT* in T cells and *NECTIN2* in fibroblasts and ESCs; *TIGIT* contains ITIM motifs in its cytoplasmic tail that binds to *NECTIN2* and triggers inhibitory signals^54^. This ligand-receptor signal was lower in P5-1 (**Fig. S5A, B**), which came from the primary tumor site. Similarly, the *IL7R_IL7* pair was observed in all four samples for fibroblast (**Fig. S5A**), with the lowest signal in P5-1 (*IL7R_IL7* was observed in P1-1 and P6-1 for ESCs, see **Fig. S5B**). This ligand-receptor pair has been correlated with immune cell infiltration in the TME^55^. The ligand *FASLG* in T cells to receptors *FAS*, *TNFRSF10A*, and *TNFRSF1B* in fibroblasts interaction pairs were detected in all samples (P5-1, 6-1 and 7-1) but not in P1-1 (**Fig. S5A**). Their interaction leads to apoptosis of thymocytes that fail to rearrange correctly their T cell receptor (TCR) genes and activation-induced cell death responsible for the peripheral deletion of activated T cells^56^. For fibroblast interactions, the *LGALS9*_*CD44/r* appear enriched in both fibroblasts and ESCs in P6-1 and P7-1, which are metastatic, with high T_inf_ (**Fig. S5A, B**). The *LGALS9*_*CD44* pair appeared on ESCs from all samples except P6-1 (**Fig. S5B**). Gal-9 has direct cytotoxic effects, binds to CD44 expressed on cancer cells to limit cancer metastasis, and enhances stability and function of adaptive regulatory T cells^57,58^. Different interactions associated with immune regulation in tumors^59^ were also found: *CD74* interactions with *APP* or *COPA* were found in samples P5-1 and P6-1, while *HLA-C_FAM3C* interaction was enriched in samples P1-1 and P7-1 (**Fig. S5A, B**). *CD2_CD58* interactions between T cells and ESCs were noted in all four samples (**Fig. S5B**), and between T cells and fibroblasts in P6-1 and P7-1 (**Fig. S5A**). *NOTCH2* interactions^60,61^ (with *JAG2* and *DLL3*) were seen in fibroblasts in P1-1 (**Fig. S5A**).

We also detected intriguing patterns of certain integrin complex-collagen binding pairs^62^ on fibroblasts enriched in specific samples (**Fig. S5C**): integrin complex *A2B1* appeared in P6-1 only; enhanced expression of α2β1 integrins may influence spheroid disaggregation and proteolysis responsible for the peritoneal dissemination of ovarian carcinoma^63^. *A1B1* was intriguingly absent in P1-1; instead integrins *A10B1* and *A11B1,* appear in P1-1 alone. Integrin α11β1^62^ was previously seen overexpressed in NSCLC, especially in CAFs^64,65^. The *CD40LG_A5B1* pair was seen for fibroblasts in P1-1, P6-1 and P7-1 (**Fig. S5A**). Integrin α5β1 plays an important role in tumor progression^66^. In addition, strong *A4B1* interactions with *FN1, VCAM1* and other ligands^67^ are seen with fibroblasts in P5-1, P6-1 and P7-1 (**Fig. S5A**), and ESCs in P6-1 (**Fig. S5B**); *A4B1* receptors have been proposed to target therapy in inflammatory disorders and cancer^67^. These results suggest that different patient samples may have unique LR signatures that are associated with specific cell types, which may be used to target therapy.

## Discussion

Ovarian cancer is a collection of different carcinomas that manifest as different histotypes, each with different cellular compositions and pathogenic mechanisms. Analysis of the TME in different ovarian cancer histotypes at the single-cell resolution can potentially connect the different histotypes with their unique cellular and molecular signatures, understand disease etiology and help guide therapy. With this aim in mind, we ran Drop-seq on 21 tumor samples from 12 patients and across 6 histotypes of ovarian cancer. We detected three major cell compartments: epithelia (epithelial cells and ESCs), stroma (fibroblast, endothelial cells, and MSCs), immune (T, B, plasma B, macrophage) by integrating all single-cell experiments. The four ovarian cancer subtypes using the TCGA gene expression signature revealed highly correlated cell types: the immunoreactive subtype showed higher correlation with immune cells, while the mesenchymal subtype correlated most with stroma cells and least with epithelial cells. The differentiated and proliferative subtypes both consisted of epithelia but with low and high percentage of ESCs, respectively. This suggests that the molecular subtypes classified by TCGA may be driven by the cell type compositions of the tumor samples. Because each tumor sample showed a unique cellular makeup that differed between primary and metastatic sites, it follows that the dominant molecular subtype of a tumor sample is specific to its site of origin, rather than being patient-specific, e.g., patients, P6 and P8, while sharing the HGSOC histotype, have different TCGA subtypes.

For most cell types, we found that the cell cycle phases G1S, G2M, and MG1 were consistently distributed with a higher percentage of MG1 phase, with the exception of ESCs, where over 70% of the ESCs belonged to the G2M phase. Tumors with high G2M gene activity have been associated with metastasis and worse outcomes in patients with particular subtypes of breast and pancreatic cancers^27,28^. The role of p53 in G2/M related cell-cycle arrest in response to DNA damage has been studied extensively^68–70^.

We found five different subtypes of cancer-associated fibroblasts, FB0 and FB2-5 in both primary and metastatic sites, based on the expression of *IL6*, *CCL2*, *S100A4*, *PDPN*, and *FGF7*. Each CAF sub-cluster supports different roles in the progression and metastasis of ovarian cancer. Cells in FB0 expressed genes associated with angiogenesis, Integrin signaling, and T cell receptor signaling pathways. These pathways were related to extracellular matrix remodeling and immune crosstalk under the tumor micro-environment (TME). FB2 supported upregulation of NF-kappa B signaling pathway genes and chemokine receptors associated with cancer metastasis. FB2 and FB4 exhibited elevated expressions of growth factor binding genes as well as genes enriched for angiogenesis and blood vessel development. Top differentially expressed genes in FB3 may be involved with endothelial cell signaling and vascular function. FB5 showed genes enriched in immune crosstalk and cytokine/interferon signaling pathways. Among epithelial cells, we identified the EP3 sub-cluster as cancer stem cells, based on high expression of *IFIT1* and *ISG15*.

The majority of the immune sub-clusters were consistent with those identified in our previous study on metastatic ovarian cancer^9^. We identified a new cluster of IL32+ B cells (BC2) that are CD38-SDC1-S100A4+GAPDH+; these cells were found in both primary and metastatic tumor sites with high T cell infiltration, deriving primarily in clear cells and SOC histotypes.

We did not observe any significant difference in the overall composition of cell lineages between primary vs. metastatic sites. We noted higher ratios of specific CAF (FB4 and FB2) and macrophage (MA2 and MA3) subsets and lower ratio of an epithelial subcluster (EP2) in metastatic sites, compared to primary tumors.

Overall percentages of T and B cells were higher in high T_Inf_ samples, be it from primary or metastatic site, while the percentage of MS was lower overall. At the sub-cluster level, the TC0 and BC3 were positively correlated with T_Inf_ status, with MS2 showing negative correlation. The CAF sub-clusters FB0, FB2, FB4 and FB5, and the CSCs (EP3) were enriched in samples classified as metastatic, low T_Inf_.

Besides tumor site and T_Inf_ status, there were also differences in the makeup of cellular sub-types between histotypes. The percentages of epithelial cells from EP1-3 were higher in HGSOC and MMMT histotypes, while the percentages of ESCs in clusters ES1-3 were higher in HGSOC only. For clear cell histotype, the percentage of cells in TC0, BC2 and BC3 was higher. The endometrioid histotype had a higher percentage of MS and FB1 cells. The percentages of MA0, MA2 and TC1 cells were higher in LGSOC than in other histotypes.

Lastly, we found fibroblasts and MSCs to be active players in the TME, exhibiting potentially distinct LR interactions with epithelial and immune subclusters in patients and histotypes. Imputed ligands and receptors may be leveraged to target therapy in ovarian cancer patients.

### Limitations of the study

The total number of patient samples collected in this study is limited due to the pandemic. Certain cancer subtypes such as MMMT were less represented in our samples because of their lower prevalence^71^. The cell sub-populations in tumors dissected from different individuals, tumor sites (primary vs. metastatic) or even different regions sampled from the same tumor may vary. The ligand-receptor interactions were inferred *in silico* through statistical testing, with the caveat that the same ligand or receptor can account for multiple inferred ligand-receptor pairs. Further validation tests are needed to confirm the ligand-receptor interactions.

## Acknowledgement

We are grateful to Drs. Ran Zhou and Preety Bajwa for helpful discussions and Dr. Mark Lingen for tissue access. All of the tissues were histologically graded, de-identified, and obtained from the University of Chicago Human Tissue Resource Center. Computational resources and data storage were provided by the University of Chicago Research Computing Center. This work was internally funded.

## AUTHOR CONTRIBUTIONS

S.O., B. X. and A.B. conceived and designed the study. S.O. performed all single-cell dissociation and scRNA-seq experiments, with support from R.B. and H.E. B.X. conducted all of the scRNA-seq data analysis. B.X., S.O., N.A.A. and A.B. interpreted the data. B.X. and A.B. wrote the manuscript, with input from S.O.

## DECLARATION OF INTERESTS

The authors declare no competing interests.

## Methods

### Tissue collection, sample preparation and Drop-seq

Ovarian cancer tissue from primary and metastatic sites were collected from women undergoing debulking surgery at the University of Chicago. Some of the tissue collected from the different sites were patient-matched. The University of Chicago’s Institutional Review Board for human research approved the collection of human tissue after patient deidentification. Ovarian tumors were transported in DMEM/F12 containing 10% FBS and 1% P/S (100% DMEMF/12) and processed as previously described^9^. Red blood cells and dead cells were removed from cell suspensions using Miltenyi Biotec, 130-094-183, 130-090-101, respectively, used according to manufacturer’s protocols. Additionally, some samples were enriched for immune-only, non-immune, tumor-only and non-tumor cell compartments, using magnetic bead-based isolation or fluorescence activated flow sorting (Miltenyi Biotec, 130-118-780, 130-045-801, 130-108-339, 130-042-401, 130-112-931, 130-118-497, 130-110-770, used according to manufacturer’s protocols).

Drop-seq was performed as previously described^9^ on ovarian cancer tumor samples from 12 patients (**Table 1**). A total of 21 tumor samples were present in this study, including 5 patients with Matched primary (right and/or left ovaries) and metastatic (omentum, rectum) tumors (**Table 2**). Of these, a few randomly selected samples were enriched for select cellular compartments prior to running Drop-seq: CD45+ (5 samples), tumor (2 samples) and non-tumor (1 sample); 18 samples were processed without any enrichment.

### Data processing, alignment and clustering analysis

A total of 40 sequencing runs were performed on Illumina’s NextSeq 500 using the 75 cycle v3 kit, as previously described^9^. Some samples were sequenced multiple times to achieve deeper resolution. Each run produced paired-end reads, with Read 1 representing the 12 bp cell barcode and a six bp long unique molecular identifier (UMI) and Read 2 representing a 60-64 bp mRNA fragment. Paired-end reads from the same samples were combined to generate 26 paired-end fastq files. Read count matrices were generated from sequence reads from the Drop-seq experiments for both exon and intron regions in the human genome (gencode^72^ hg38 v.27) using a *Snakemake* pipeline^73^ and STAR version 2.5.3 aligner^74^.

To select high-quality cells, we applied a filter based on the number of genes detected per cell. Prior to filtering, each sequenced sample produced approximately 5,000 cells. Based on the median number of captured genes per cell, cells with less than 400 genes detected were removed from the dataset. A total of 26,421 cells were retained for the downstream analysis. We followed a standardized pipeline using the single-cell analysis tool suite, Seurat v3.0.2^75,76^. A logarithmic normalization method^75^ was applied to all samples to transfer the gene expression counts (+1, to avoid log(0)) scaled by a factor of 10,000 (TP10K) to log units. The normalized matrices for all samples were integrated by the anchor-based alignment method Canonical Correlation Analysis (CCA) using Seurat^76^. The top 1,311 highly variable genes and top 20 canonical vectors were selected to perform the alignment integration, where the integrated gene expression matrix had a lesser number of features (genes) than the original gene expression matrix. The integrated matrix was scaled by a linear transformation to center the mean gene expression for all cells. We applied PCA on the scaled integrated expression matrix to extract the top 50 components in the data, followed by a heuristic elbow plot on the standard deviation of each PC. We selected the top 16 variant PCs based on the elbow plot. The selected PCs were used in further exploration of the data, such as UMAP^77^ dimension reduction, construction of K-nearest neighbor graphs, shared nearest neighbor modularity optimization-based clustering^76^, etc. We used dimension reduction methods, UMAP, to generate 2D plots to visualize different cell populations in the experiments. Hierarchical clustering on the shared nearest neighbor graph was applied to infer the clustering structure on the cells where the resolution parameter was set to 0.2. Differential expression analysis was performed through the *FindMarkers* function in Seurat using the Wilcoxon Rank Sum test, and statistically significant markers were extracted for sub-populations or contrast groups based on an adjusted p-value (adj. p-val.) threshold of 0.05.

### Cell cycling effects

We inferred the cell-cycle phase for all cells based on previously curated gene markers reflecting three phases of the cell cycle in chemically synchronized cells (G1/S, G2/M and M/G1)^5,78^. For each cell-cycle phase, the module scores were calculated as the average expression levels of binned gene markers subtracted by the aggregated expression of random gene sets from the same bin. The Seurat *AddModuleScore* function was used to assign all five module scores to each cell where 24 bins of aggregate expression levels for the marker genes were used and a hundred control genes were selected from the same bin per gene. The highest scored cell-cycle phase was assigned to the cell. If none of the module scores were positive, the cell was designated as not assigned (NA).

### Cancer subtype classification

Four cancer subtypes-differentiated, immunoreactive, mesenchymal, and proliferative were categorized by previous bulk sequencing study in ovarian cancer^3,79^. The marker genes for each subtype were determined by the upregulated marker signatures on the four subtypes^80^. The Seurat *AddModuleScore* function was used to assign four module scores to each cell where 24 bins of aggregate expression levels for the marker genes were used and a hundred control genes were selected from the same bin per gene. The subtypes were then assigned to individual cells by the highest positive modular score. In the absence of positive modular scores, the subtype was considered not assigned (NA).

### Cancer patient survival prediction

The cancer outcome was categorized as poor and good in the previous research on the TCGA ovarian cancer dataset^3,79^, where a list of gene signatures based on RNA-seq data were extracted for both outcomes. We obtained the module scores based on these lists of predictive gene markers using the Seurat *AddModuleScore* function as described in the cancer subtype classification. The predicted outcome was assigned to the cells according to the module score.

### Cell type classification using template-based method

We assigned the cell type using a template-based cell annotation method, namely sc-TACA (https://github.com/bingqing-Xie/taca)^81^. The sc-TACA method utilizes annotated single-cell dataset as a template. In this study, six HGSOC metastatic samples in the 26 samples have been previously annotated, which was used as the template. The cell types annotated in this template were denoted by *T* = {*t*_i_, *t* = 1. . *p*}, where *p* is the total number of unique cell types. All samples were integrated by an anchor-based alignment via Canonical Correlation Analysis (CCA) in Seurat^75,76^. Then modularity optimization-based hierarchical clustering *FindClusters* was applied on the integrated dataset with a resolution r = 0.2 that resulted in 11 cell clusters. For each cluster *i*, we obtained the annotated cell type vector *C*_i_ = {*c*_l_, *c*_2_, …, *c*_Ni_} where *N*_i_ is the total number of cells from cluster *i* and *c*_i_ ∈ *T*. The annotation *t*_i_ of a given cluster *i* was determined by highest ratio of annotated cell type within the cluster 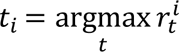 where 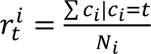. A threshold *r*_min_ = 0.7 was enforced to ensure the robust assignment. If 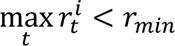 for cluster *i*, it was labeled as undecided.

### Immunohistochemistry

Ovarian cancer tissues were fixed and stained for Immunohistochemistry as previously described^9^, to evaluate the fraction of cytokeratin-7 (Thermo Scientific, MA5-11986), pan-vimentin (Abcam, Ab16700), CD45 (Agilent, M0701) positive cells.

Aperio ImageScope v12.4.3 was used to analyze the fraction of cells that stained for CD45, vimentin and CK7 in the entire tissue section, using algorithm ‘Positive Pixel Count v9’.

### Analysis on fibroblasts, epithelial cells, and immune sub-population

After identifying the cell types, we extracted the fibroblasts, epithelial cells, and immune cells (T cells, B cells, macrophages) to conduct further investigation. Each sub-population expression matrix was subset from the integrated matrix. The expression matrix was scaled and PCA analysis was performed to extract top components in the data. Top PCs were selected based on the elbow plot, which varied from 10 to 20 based on the sub-population variation. Hierarchical clustering on the shared nearest neighbor graph was applied where the resolution parameter was set to a range between 0.2 and 0.5. The same UMAP was used to project the cells to a 2D space to visualize the sub-types for each cell type. Differential expression analysis was performed through the *FindMarkers* function in Seurat using the Wilcoxon Rank Sum test, and statistically significant markers were extracted for sub-populations or contrast groups based on an adjusted p-value (adj. p-val.) threshold of 0.05. The differences in cell composition ratios between primary and metastatic sites, and between high and low T_inf_ groups were evaluated by two-sided T-test with P value estimation.

### Ligand-receptor interaction analysis on cell type subclusters

We constructed a customized pan-cancer ligand-receptor (LR) interaction database, using *CellphoneDB*^53^ and published cancer studies, including 27 immune checkpoint LR pairs^82^, 114 interaction pairs between cancer cells and T cells in lung cancer^83^, 1380 LR pairs in a pan-cancer study^84^, and 216 LR pairs related to ovarian cancer^85^. For each sample, we inferred LR interactions among any pair of the cell sub-clusters, 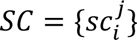, where *i* is the lineage such as EP, and *j* is the subcluster index, using the pan-cancer LR database. We obtained a P value for the likelihood of cell-type enrichment of each ligand– receptor complex (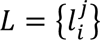, where *i* is a ligand and *j* is a corresponding receptor). We denote 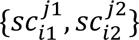 for a sub-clusters pair. P value is calculated by the proportion of means that are as high as (or higher) than the random permutation for all pairs, *SC* = 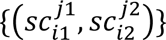. Interactions with adjusted P value < 0.05 were considered significant. The ‘significant means’ vector, 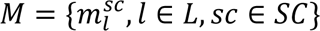 was extracted for each sample and *m*_l_ was set to 0 when P value > 0.05 or sub-clusters with insufficient cell counts, (|*sc*|) < 50. The number of absolute interactions ∑(*M* > 0) was used as a proxy to estimate the frequency of the cell-cell crosstalk among cell types. The hierarchical clustered heatmap was used to identify the shared patterns for the sub-clusters from different cell types. We then grouped the samples by histotype and site for the downstream comparative analysis. A linear model was built using *lmfit* in Limma R library for a given contrast group (e.g. one histotype against the rest of the histotypes) and the empirical Bayes moderated t-statistics test *ebayes* was used to estimate the significance of any LR signature, *l* ∈ *L*^86,87^. Significant positive LR pairs were used as the signature for any given condition group.

### Data Availability

The data discussed in this publication have been deposited in NCBI’s Gene Expression Omnibus^88^ and are accessible through GEO Series accession number GSE235931 (https://www.ncbi.nlm.nih.gov/geo/query/acc.cgi?acc=GSE235931).

## Notes

https://www.ncbi.nlm.nih.gov/geo/query/acc.cgi?acc=GSE235931

### Competing Interest Statement

The authors have declared no competing interest.

## References

1 Matulonis, U. A. et al. Ovarian cancer. Nature Reviews Disease Primers 2, 16061, doi:10.1038/nrdp.2016.61 (2016).

2 Society, A. American Cancer Society. Cancer Facts & Figures 2020. Am Cancer Soc, 1-52 (2020).

3 Network, C. G. A. R. Integrated genomic analyses of ovarian carcinoma. Nature 474, 609 (2011).

4 Sangha, N. et al. Neurofibromin 1 (NF1) defects are common in human ovarian serous carcinomas and co-occur with TP53 mutations. Neoplasia 10, 1362–1372, following 1372, doi:10.1593/neo.08784 (2008).

5 Evan et al. Highly Parallel Genome-wide Expression Profiling of Individual Cells Using Nanoliter Droplets. Cell 161, 1202–1214, doi:10.1016/j.cell.2015.05.002 (2015).

6 Izar, B. et al. A single-cell landscape of high-grade serous ovarian cancer. Nature Medicine 26, 1271–1279, doi:10.1038/s41591-020-0926-0 (2020).

7 Zhang, K. et al. Longitudinal single-cell RNA-seq analysis reveals stress-promoted chemoresistance in metastatic ovarian cancer. Science Advances 8, eabm1831, doi:doi:10.1126/sciadv.abm1831 (2022).

8 Kan, T. et al. Single-cell RNA-seq recognized the initiator of epithelial ovarian cancer recurrence. Oncogene 41, 895-906, doi:10.1038/s41388-021-02139-z (2022).

9 Olalekan, S., Xie, B., Back, R., Eckart, H. & Basu, A. Characterizing the tumor microenvironment of metastatic ovarian cancer by single-cell transcriptomics. Cell Reports 35, 109165, doi:10.1016/j.celrep.2021.109165 (2021).

10 Konecny, G. E. et al. Prognostic and Therapeutic Relevance of Molecular Subtypes in High-Grade Serous Ovarian Cancer. JNCI: Journal of the National Cancer Institute 106, dju249–dju249, doi:10.1093/jnci/dju249 (2014).

11 Tay, D. L., Bhathal, P. S. & Fox, R. M. Quantitation of G0 and G1 phase cells in primary carcinomas. Antibody to M1 subunit of ribonucleotide reductase shows G1 phase restriction point block. Journal of Clinical Investigation 87, 519–527, doi:10.1172/jci115026 (1991).

12 Becker, K. A. et al. Self-renewal of human embryonic stem cells is supported by a shortened G1 cell cycle phase. Journal of Cellular Physiology 209, 883–893, doi:10.1002/jcp.20776 (2006).

13 Aass, K. R. et al. Intracellular IL-32 regulates mitochondrial metabolism, proliferation, and differentiation of malignant plasma cells. iScience 25, 103605, doi:10.1016/j.isci.2021.103605 (2022).

14 Hu, Z. et al. The Repertoire of Serous Ovarian Cancer Non-genetic Heterogeneity Revealed by Single-Cell Sequencing of Normal Fallopian Tube Epithelial Cells. Cancer Cell 37, 226–242.e227, doi:10.1016/j.ccell.2020.01.003 (2020).

15 Schneider, M., Hansen, J. L. & Sheikh, S. P. S100A4: a common mediator of epithelial– mesenchymal transition, fibrosis and regeneration in diseases? Journal of Molecular Medicine 86, 507–522, doi:10.1007/s00109-007-0301-3 (2008).

16 Cui, Y. et al. Dual effects of targeting S100A11 on suppressing cellular metastatic properties and sensitizing drug response in gastric cancer. Cancer Cell International 21, doi:10.1186/s12935-021-01949-1 (2021).

17 Wang, D., You, D. & Li, L. Galectin-3 regulates chemotherapy sensitivity in epithelial ovarian carcinoma via regulating mitochondrial function. The Journal of Toxicological Sciences 44, 47–56, doi:10.2131/jts.44.47 (2019).

18 Kim, J. et al. High-grade serous ovarian cancer arises from fallopian tube in a mouse model. Proceedings of the National Academy of Sciences 109, 3921–3926, doi:10.1073/pnas.1117135109 (2012).

19 Scurr, L. L. et al. Ankyrin Repeat Domain 1, ANKRD1, a Novel Determinant of Cisplatin Sensitivity Expressed in Ovarian Cancer. Clinical Cancer Research 14, 6924–6932, doi:10.1158/1078-0432.ccr-07-5189 (2008).

20 Jen, J. & Wang, Y.-C. Zinc finger proteins in cancer progression. Journal of Biomedical Science 23, doi:10.1186/s12929-016-0269-9 (2016).

21 Behera, A., Ashraf, R., Srivastava, A.K., Kumar, S. Bioinformatics analysis and verification of molecular targets in ovarian cancer stem-like cells. Heliyon 6 (2020).

22 Hu, Y. et al. Identification of a five-gene signature of the RGS gene family with prognostic value in ovarian cancer. Genomics 113, 2134–2144, doi:10.1016/j.ygeno.2021.04.012 (2021).

23 Barber, E. K., Dasgupta, J. D., Schlossman, S. F., Trevillyan, J. M. & Rudd, C. E. The CD4 and CD8 antigens are coupled to a protein-tyrosine kinase (p56lck) that phosphorylates the CD3 complex. Proceedings of the National Academy of Sciences 86, 3277–3281, doi:10.1073/pnas.86.9.3277 (1989).

24 Li, W. et al. Unraveling the roles of CD44/CD24 and ALDH1 as cancer stem cell markers in tumorigenesis and metastasis. Scientific Reports 7, doi:10.1038/s41598-017-14364-2 (2017).

25 Zhou, X., Hu, W. & Qin, X. The Role of Complement in the Mechanism of Action of Rituximab for B-Cell Lymphoma: Implications for Therapy. The Oncologist 13, 954–966, doi:10.1634/theoncologist.2008-0089 (2008).

26 Chen, J. et al. CD59 Regulation by SOX2 Is Required for Epithelial Cancer Stem Cells to Evade Complement Surveillance. Stem Cell Reports 8, 140–151, doi:10.1016/j.stemcr.2016.11.008 (2017).

27 Oshi, M. et al. High G2M Pathway Score Pancreatic Cancer is Associated with Worse Survival, Particularly after Margin-Positive (R1 or R2) Resection. Cancers 12, 2871 (2020).

28 Oshi, M. et al. G2M Cell Cycle Pathway Score as a Prognostic Biomarker of Metastasis in Estrogen Receptor (ER)-Positive Breast Cancer. International Journal of Molecular Sciences 21, 2921 (2020).

29 Jung, Y., Kraikivski, P., Shafiekhani, S., Terhune, S. S. & Dash, R. K. Crosstalk between Plk1, p53, cell cycle, and G2/M DNA damage checkpoint regulation in cancer: computational modeling and analysis. NPJ Syst Biol Appl 7, 46, doi:10.1038/s41540-021-00203-8 (2021).

30 Mirzayans, R., Andrais, B., Scott, A. & Murray, D. New insights into p53 signaling and cancer cell response to DNA damage: implications for cancer therapy. J Biomed Biotechnol 2012, 170325, doi:10.1155/2012/170325 (2012).

31 Viner-Breuer, R., Yilmaz, A., Benvenisty, N. & Goldberg, M. The essentiality landscape of cell cycle related genes in human pluripotent and cancer cells. Cell Division 14, 15, doi:10.1186/s13008-019-0058-4 (2019).

32 Taylor, W. R. & Stark, G. R. Regulation of the G2/M transition by p53. Oncogene 20, 1803–1815, doi:10.1038/sj.onc.1204252 (2001).

33 Kalluri, R. The biology and function of fibroblasts in cancer. Nature Reviews Cancer 16, 582–598, doi:10.1038/nrc.2016.73 (2016).

34 Liu, T., Zhou, L., Li, D., Andl, T. & Zhang, Y. Cancer-Associated Fibroblasts Build and Secure the Tumor Microenvironment. Frontiers in Cell and Developmental Biology 7, doi:10.3389/fcell.2019.00060 (2019).

35 Puram, S. V. et al. Single-Cell Transcriptomic Analysis of Primary and Metastatic Tumor Ecosystems in Head and Neck Cancer. Cell 171, 1611–1624.e1624, doi:10.1016/j.cell.2017.10.044 (2017).

36 Neri, S. et al. Podoplanin-expressing cancer-associated fibroblasts lead and enhance the local invasion of cancer cells in lung adenocarcinoma. International Journal of Cancer 137, 784–796, doi:10.1002/ijc.29464 (2015).

37 Ye, J. et al. Increased expression of immediate early response gene 3 protein promotes aggressive progression and predicts poor prognosis in human bladder cancer. BMC Urology 18, doi:10.1186/s12894-018-0388-6 (2018).

38 Sang, Y. et al. SGK1 in Human Cancer: Emerging Roles and Mechanisms. Frontiers in Oncology 10, doi:10.3389/fonc.2020.608722 (2021).

39 Yang, Y., Xin, X., Fu, X. & Xu, D. Expression pattern of human SERPINE2 in a variety of human tumors. Oncology Letters, doi:10.3892/ol.2018.7819 (2018).

40 Provance, O. K. & Lewis-Wambi, J. Deciphering the role of interferon alpha signaling and microenvironment crosstalk in inflammatory breast cancer. Breast Cancer Research 21, doi:10.1186/s13058-019-1140-1 (2019).

41 Middleton, J. et al. A comparative study of endothelial cell markers expressed in chronically inflamed human tissues: MECA-79, Duffy antigen receptor for chemokines, von Willebrand factor, CD31, CD34, CD105 and CD146. The Journal of Pathology 206, 260–268, doi:10.1002/path.1788 (2005).

42 Liu, Z. et al. ENDOGLIN Is Dispensable for Vasculogenesis, but Required for Vascular Endothelial Growth Factor-Induced Angiogenesis. PLoS ONE 9, e86273, doi:10.1371/journal.pone.0086273 (2014).

43 Bao, L. et al. The S100A6 calcium-binding protein regulates endothelial cell-cycle progression and senescence. FEBS Journal 279, 4576–4588, doi:10.1111/febs.12044 (2012).

44 Salazar, N. a. B. A. Z. Support of Tumor Endothelial Cells by Chemokine Receptors. Frontiers in immunology 10 (2019).

45 Lee, J. S. et al. Duffy Antigen Facilitates Movement of Chemokine Across the Endothelium In Vitro and Promotes Neutrophil Transmigration In Vitro and In Vivo. The Journal of Immunology 170, 5244–5251, doi:10.4049/jimmunol.170.10.5244 (2003).

46 Peiper, S. C. et al. The Duffy antigen/receptor for chemokines (DARC) is expressed in endothelial cells of Duffy negative individuals who lack the erythrocyte receptor. Journal of Experimental Medicine 181, 1311–1317, doi:10.1084/jem.181.4.1311 (1995).

47 Berns, H., Humar, R., Hengerer, B., Kiefer, F. N. & Battegay, E. J. RACK1 IS UP-REGULATED IN ANGIOGENESIS AND HUMAN CARCINOMAS. The FASEB Journal 14, 2549–2558, doi:10.1096/fj.99-1038com (2000).

48 Maharshak, N. CD74 is a survival receptor on colon epithelial cells. World Journal of Gastroenterology 16, 3258, doi:10.3748/wjg.v16.i26.3258 (2010).

49 Köbel, M. et al. An Immunohistochemical Algorithm for Ovarian Carcinoma Typing. Int J Gynecol Pathol 35, 430–441, doi:10.1097/pgp.0000000000000274 (2016).

50 Peres, L. C. et al. Histotype classification of ovarian carcinoma: A comparison of approaches. Gynecol Oncol 151, 53–60, doi:10.1016/j.ygyno.2018.08.016 (2018).

51 Leskela, S. et al. Molecular Basis of Tumor Heterogeneity in Endometrial Carcinosarcoma. Cancers (Basel) 11, doi:10.3390/cancers11070964 (2019).

52 Sakata, J. et al. Impact of positive ZEB1 expression in patients with epithelial ovarian carcinoma as an oncologic outcome-predicting indicator. Oncol Lett 14, 4287–4293, doi:10.3892/ol.2017.6658 (2017).

53 Efremova, M., Vento-Tormo, M., Teichmann, S. A. & Vento-Tormo, R. CellPhoneDB: inferring cell–cell communication from combined expression of multi-subunit ligand–receptor complexes. Nature Protocols 15, 1484–1506, doi:10.1038/s41596-020-0292-x (2020).

54 Deuss, F. A., Gully, B. S., Rossjohn, J. & Berry, R. Recognition of nectin-2 by the natural killer cell receptor T cell immunoglobulin and ITIM domain (TIGIT). Journal of Biological Chemistry 292, 11413–11422, doi:10.1074/jbc.m117.786483 (2017).

55 Wang, X. et al. IL7R Is Correlated With Immune Cell Infiltration in the Tumor Microenvironment of Lung Adenocarcinoma. Front Pharmacol 13, 857289, doi:10.3389/fphar.2022.857289 (2022).

56 Volpe, E., Sambucci, M., Battistini, L. & Borsellino, G. Fas-Fas Ligand: Checkpoint of T Cell Functions in Multiple Sclerosis. Front Immunol 7, 382, doi:10.3389/fimmu.2016.00382 (2016).

57 Ustyanovska Avtenyuk, N., et al. Galectin-9 Triggers Neutrophil-Mediated Anticancer Immunity. Biomedicines 10, doi:10.3390/biomedicines10010066 (2021).

58 Wu, C. et al. Galectin-9-CD44 Interaction Enhances Stability and Function of Adaptive Regulatory T Cells. Immunity 41, 270–282, doi:10.1016/j.immuni.2014.06.011 (2014).

59 Li, C., Guo, L., Li, S. & Hua, K. Single-cell transcriptomics reveals the landscape of intra-tumoral heterogeneity and transcriptional activities of ECs in CC. Molecular Therapy-Nucleic Acids 24, 682–694 (2021).

60 Galic, V. et al. NOTCH2 expression is decreased in epithelial ovarian cancer and is related to the tumor histological subtype. Pathology discovery 1, 4 (2013).

61 Jia, D., Underwood, J., Xu, Q. & Xie, Q. NOTCH2/NOTCH3/DLL3/MAML1/ADAM17 signaling network is associated with ovarian cancer. Oncology Letters 17, 4914–4920 (2019).

62 Zeltz, C. & Gullberg, D. The integrin–collagen connection–a glue for tissue repair? Journal of cell science 129, 653–664 (2016).

63 Shield, K. et al. Alpha2beta1 integrin affects metastatic potential of ovarian carcinoma spheroids by supporting disaggregation and proteolysis. J Carcinog 6, 11, doi:10.1186/1477-3163-6-11 (2007).

64 Navab, R. et al. Prognostic gene-expression signature of carcinoma-associated fibroblasts in non-small cell lung cancer. Proc Natl Acad Sci U S A 108, 7160–7165, doi:10.1073/pnas.1014506108 (2011).

65 Zhu, C.-Q. et al. Integrin α11 regulates IGF2 expression in fibroblasts to enhance tumorigenicity of human non-small-cell lung cancer cells. Proceedings of the National Academy of Sciences 104, 11754–11759, doi:doi:10.1073/pnas.0703040104 (2007).

66 Hou, J., Yan, D., Liu, Y., Huang, P. & Cui, H. The roles of integrin α5β1 in human cancer. OncoTargets and therapy 13, 13329 (2020).

67 Baiula, M., Spampinato, S., Gentilucci, L. & Tolomelli, A. Novel ligands targeting α4β1 integrin: Therapeutic applications and perspectives. Frontiers in Chemistry 7, 489 (2019).

68 Concin, N. et al. Role of p53 in G2/M cell cycle arrest and apoptosis in response to gamma-irradiation in ovarian carcinoma cell lines. Int J Oncol 22, 51–57 (2003).

69 Agarwal, M. L., Agarwal, A., Taylor, W. R. & Stark, G. R. p53 controls both the G2/M and the G1 cell cycle checkpoints and mediates reversible growth arrest in human fibroblasts. Proc Natl Acad Sci U S A 92, 8493–8497, doi:10.1073/pnas.92.18.8493 (1995).

70 Müller, I. et al. Cancer Cells Employ Nuclear Caspase-8 to Overcome the p53-Dependent G2/M Checkpoint through Cleavage of USP28. Molecular Cell 77, 970–984.e977, doi:10.1016/j.molcel.2019.12.023 (2020).

71 Xu, Q., Zhang, X. & Zou, Y. Primitive ovarian carcinosarcoma: a clinical and radiological analysis of five cases. Journal of Ovarian Research 13, 1–8 (2020).

72 Frankish, A. et al. GENCODE reference annotation for the human and mouse genomes. Nucleic Acids Research 47, D766–D773, doi:10.1093/nar/gky955 (2019).

73 Selewa, A. et al. Systematic Comparison of High-throughput Single-Cell and Single-Nucleus Transcriptomes during Cardiomyocyte Differentiation. Scientific Reports 10, doi:10.1038/s41598-020-58327-6 (2020).

74 Dobin, A. et al. STAR: ultrafast universal RNA-seq aligner. Bioinformatics 29, 15–21, doi:10.1093/bioinformatics/bts635 (2013).

75 Butler, A., Hoffman, P., Smibert, P., Papalexi, E. & Satija, R. Integrating single-cell transcriptomic data across different conditions, technologies, and species. Nature Biotechnology 36, 411, doi:10.1038/nbt.4096 (2018).

76 Stuart, T. et al. Comprehensive Integration of Single-Cell Data. Cell 177, 1888–1902.e1821, doi:10.1016/j.cell.2019.05.031 (2019).

77 McInnes, L., Healy, J. & Melville, J. UMAP: Uniform Manifold Approximation and Projection for Dimension Reduction. Journal of Open Source Software 29, 861, doi:10.21105/joss.00861 (2018).

78 Whitfield, M. L. et al. Identification of Genes Periodically Expressed in the Human Cell Cycle and Their Expression in Tumors. Molecular Biology of the Cell 13, 1977–2000, doi:10.1091/mbc.02-02-0030 (2002).

79 Tothill, R. W. et al. Novel Molecular Subtypes of Serous and Endometrioid Ovarian Cancer Linked to Clinical Outcome. Clinical Cancer Research 14, 5198–5208, doi:10.1158/1078-0432.ccr-08-0196 (2008).

80 Verhaak, R. G. W. et al. Prognostically relevant gene signatures of high-grade serous ovarian carcinoma. Journal of Clinical Investigation, doi:10.1172/jci65833 (2012).

81 Xie, B. Template-based automatic cell annotation (TACA), <https://github.com/bingqing-Xie/taca> (2021).

82 Pardoll, D. M. The blockade of immune checkpoints in cancer immunotherapy. Nature Reviews Cancer 12, 252–264, doi:10.1038/nrc3239 (2012).

83 Chen, Z. et al. Ligand-receptor interaction atlas within and between tumor cells and T cells in lung adenocarcinoma. International Journal of Biological Sciences 16, 2205–2219, doi:10.7150/ijbs.42080 (2020).

84 Ghoshdastider, U. et al. Pan-Cancer Analysis of Ligand–Receptor Cross-talk in the Tumor Microenvironment. Cancer Research 81, 1802–1812, doi:10.1158/0008-5472.can-20-2352 (2021).

85 Castellano, G. et al. New Potential Ligand-Receptor Signaling Loops in Ovarian Cancer Identified in Multiple Gene Expression Studies. Cancer Research 66, 10709–10719, doi:10.1158/0008-5472.can-06-1327 (2006).

86 Dixon, P. VEGAN, a package of R functions for community ecology. Journal of Vegetation Science 14, 927–930, doi:10.1111/j.1654-1103.2003.tb02228.x (2003).

87 Smyth, G. K. in Bioinformatics and computational biology solutions using R and Bioconductor 397–420 (Springer-Verlag, 2005).

88 Edgar, R., Domrachev, M. & Lash, A. E. Gene Expression Omnibus: NCBI gene expression and hybridization array data repository. Nucleic Acids Res 30, 207–210, doi:10.1093/nar/30.1.207 (2002).

