## Supplementary figures and images for "Exploring the tumor micro-environment in ovarian cancer histotypes and tumor sites"

### Supplemental Figure 1

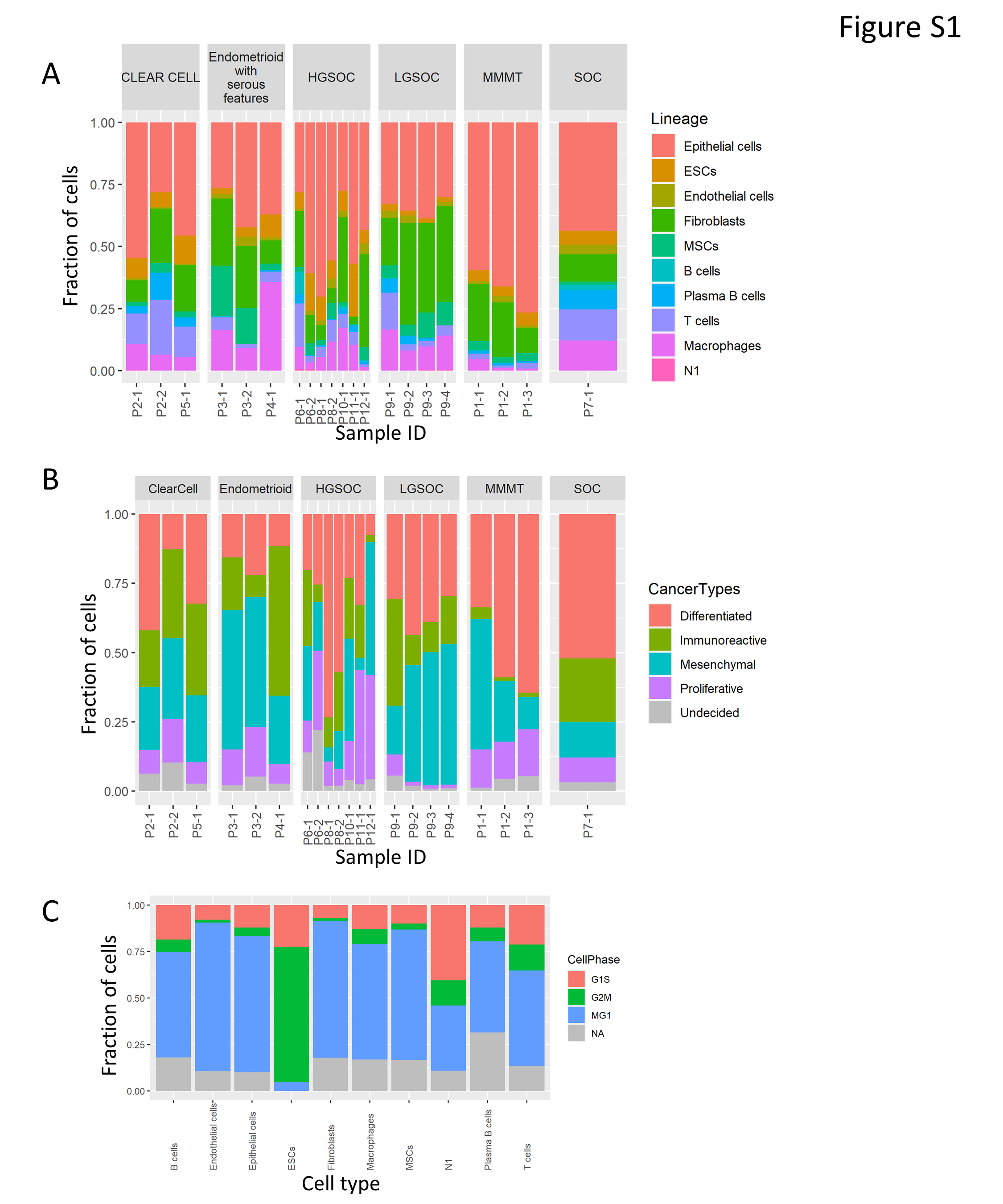

### Supplemental Figure 2

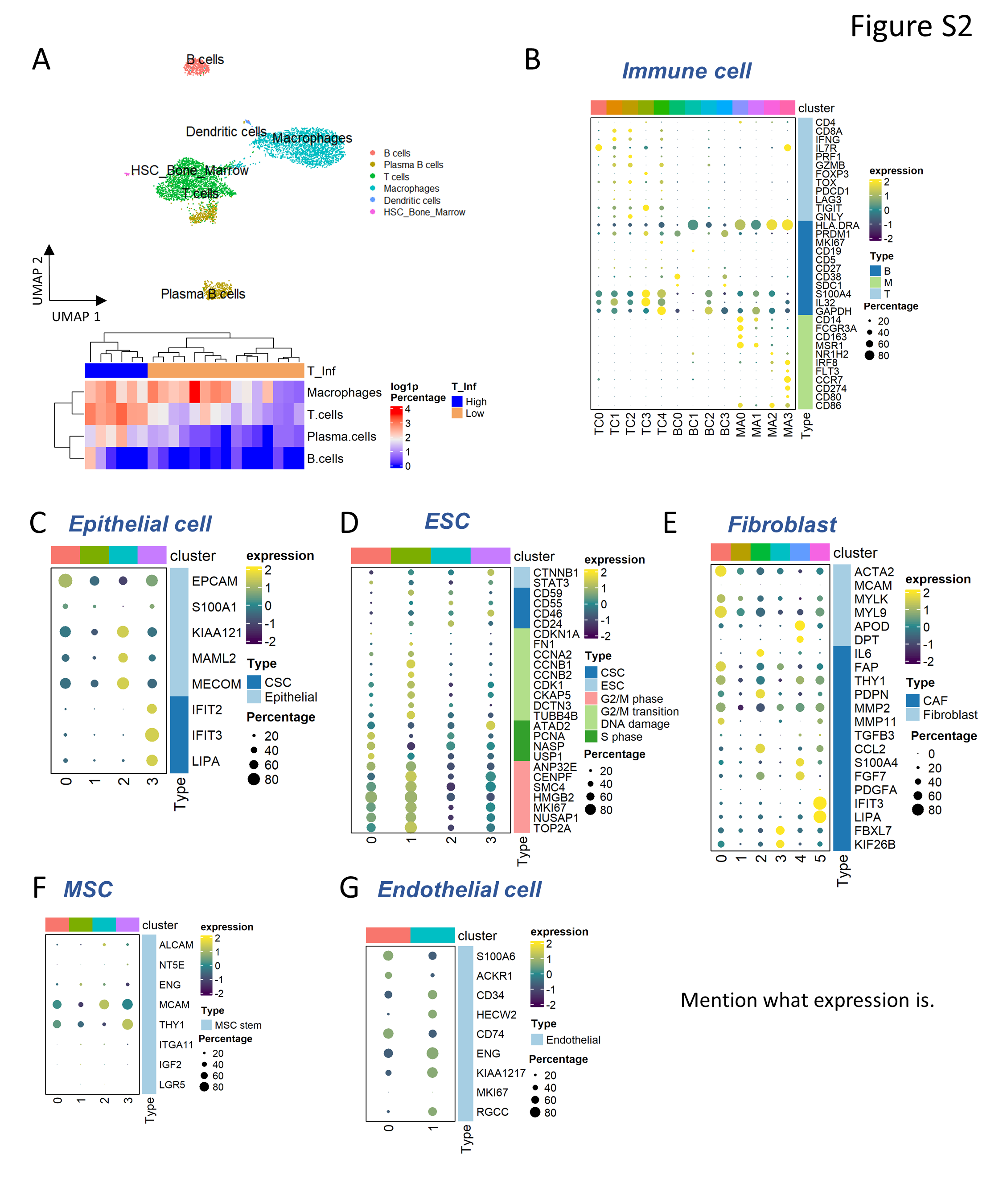

### Supplemental Figure 3

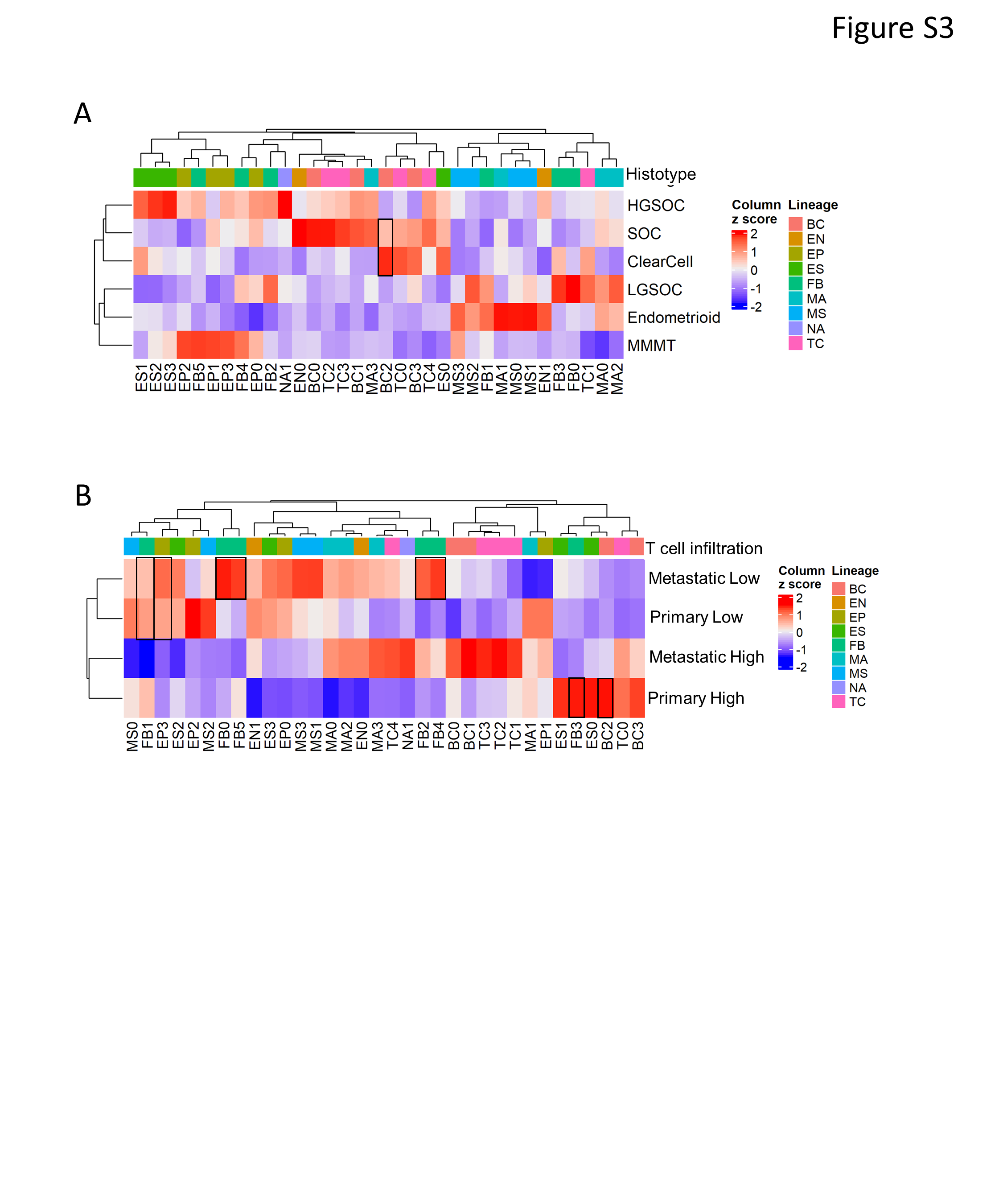

### Supplemental Figure 3C

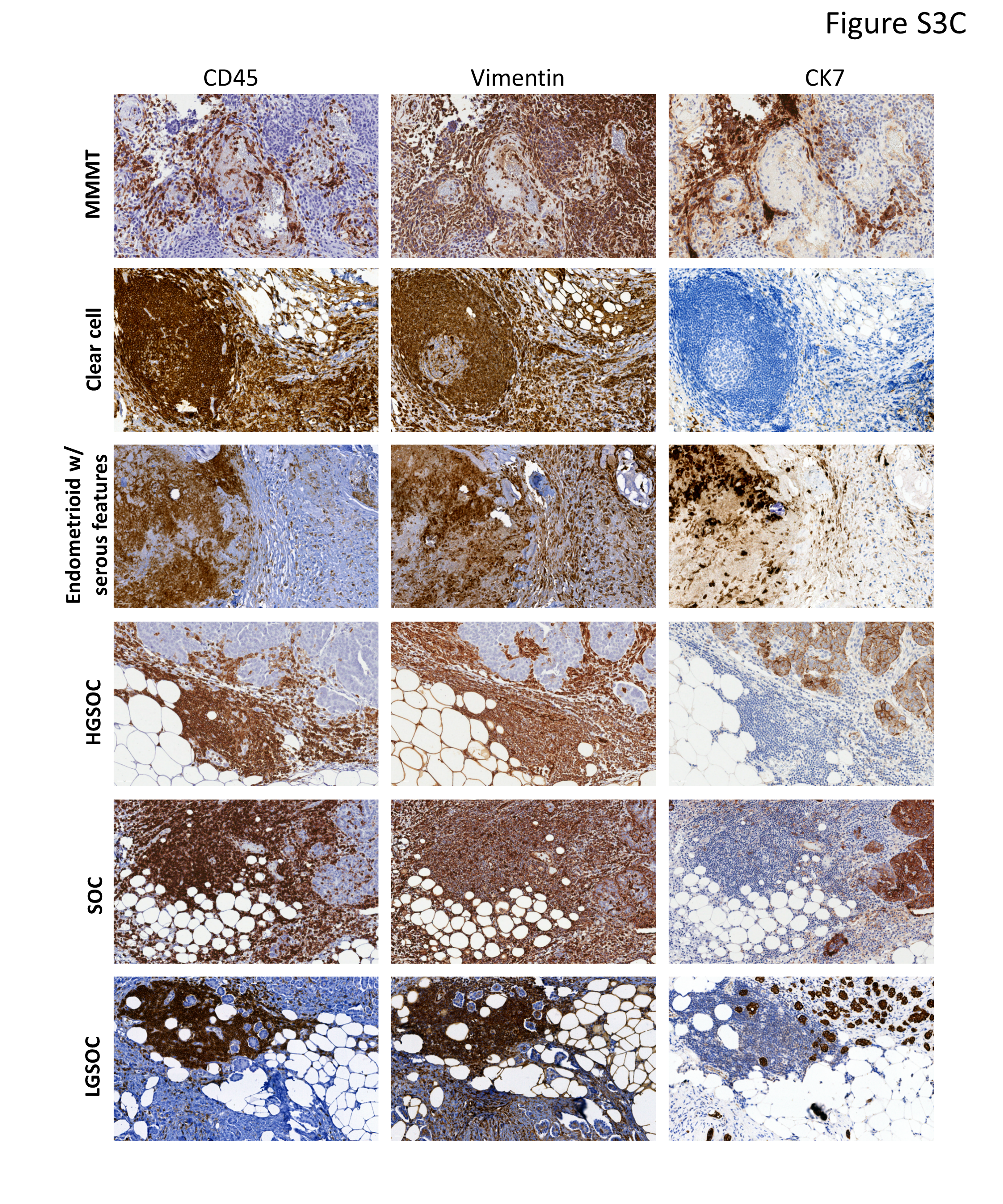

### Supplemental Figure 3D

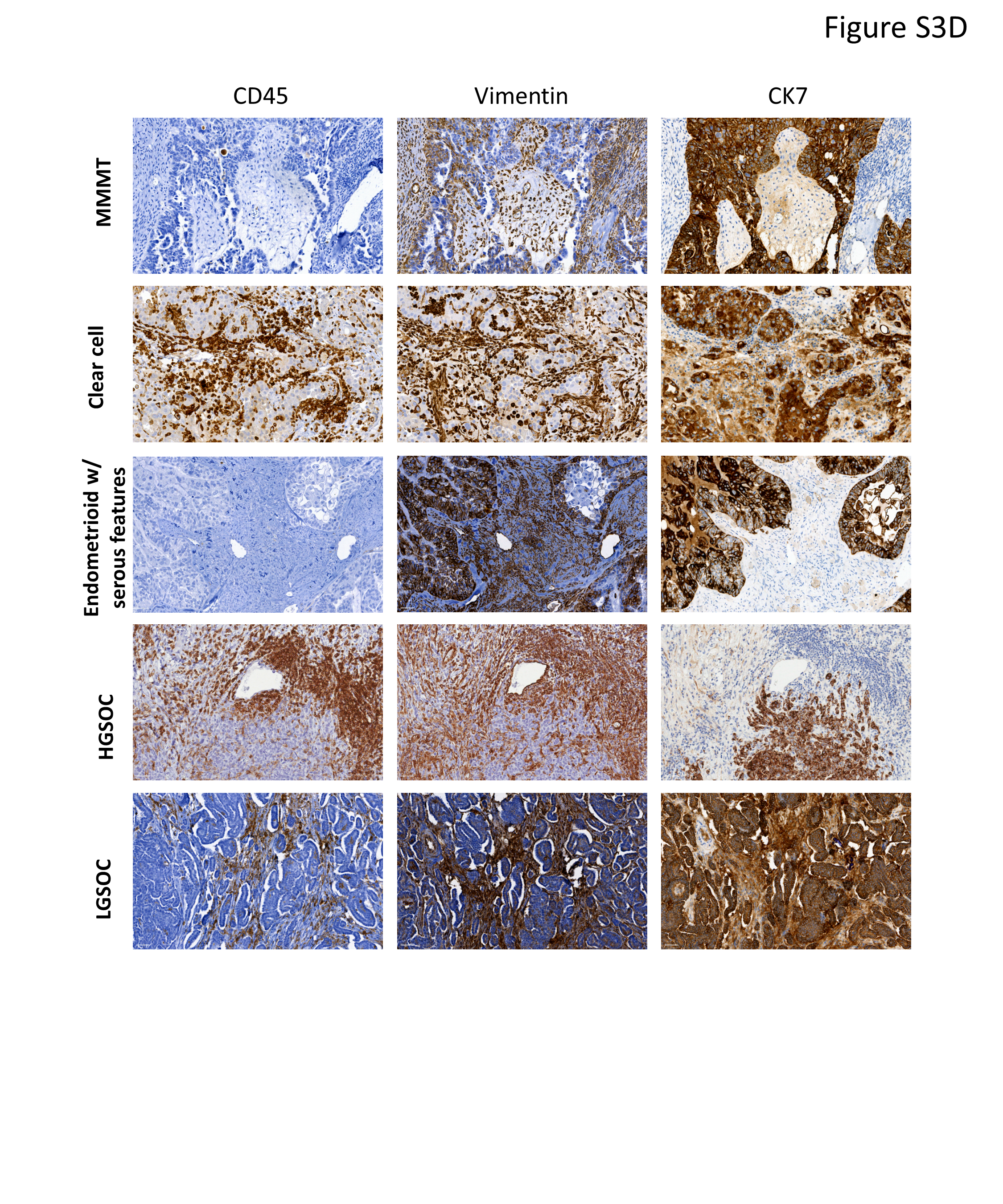

### Supplemental Figure 4A

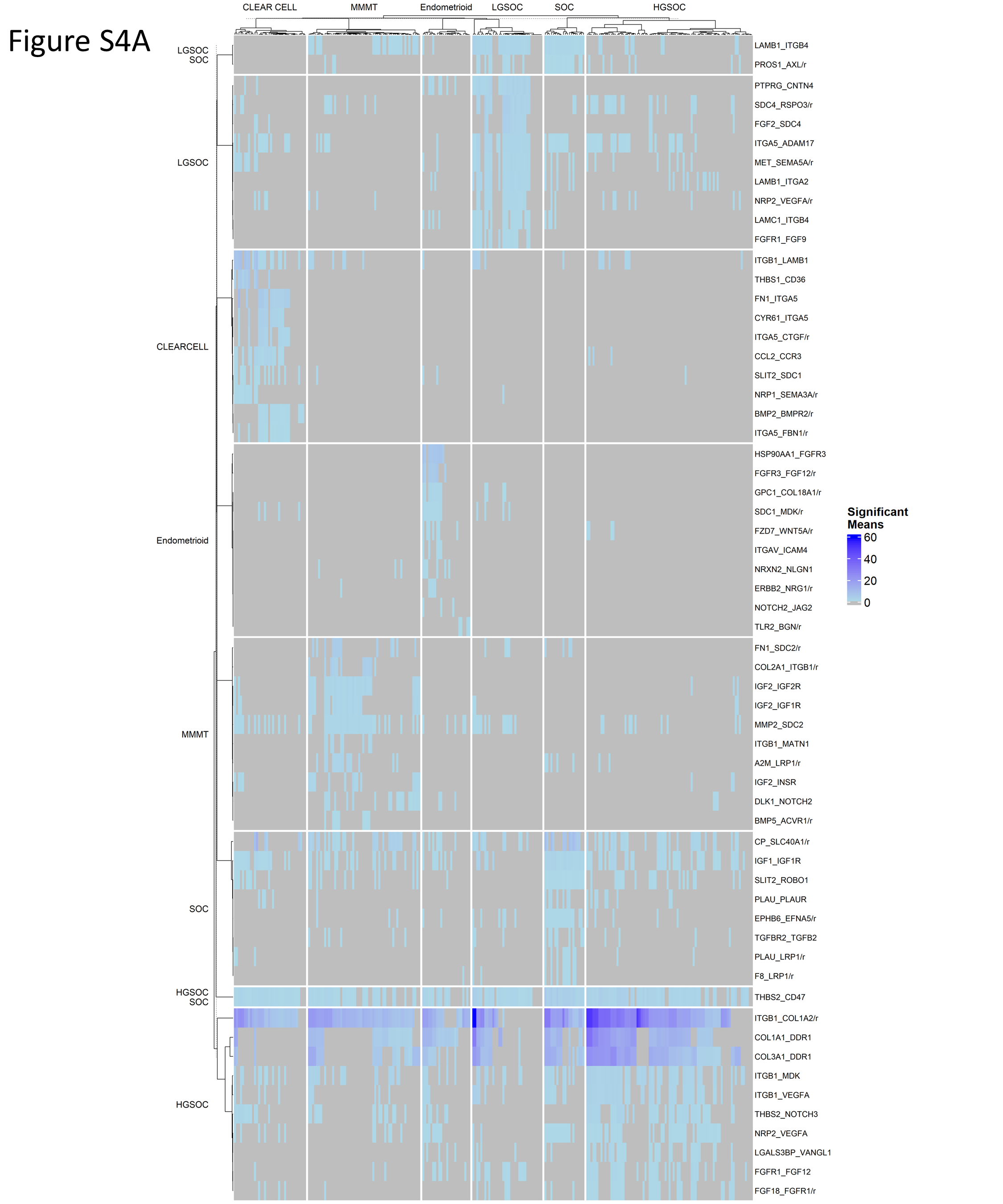

### Supplemental Figure 4B

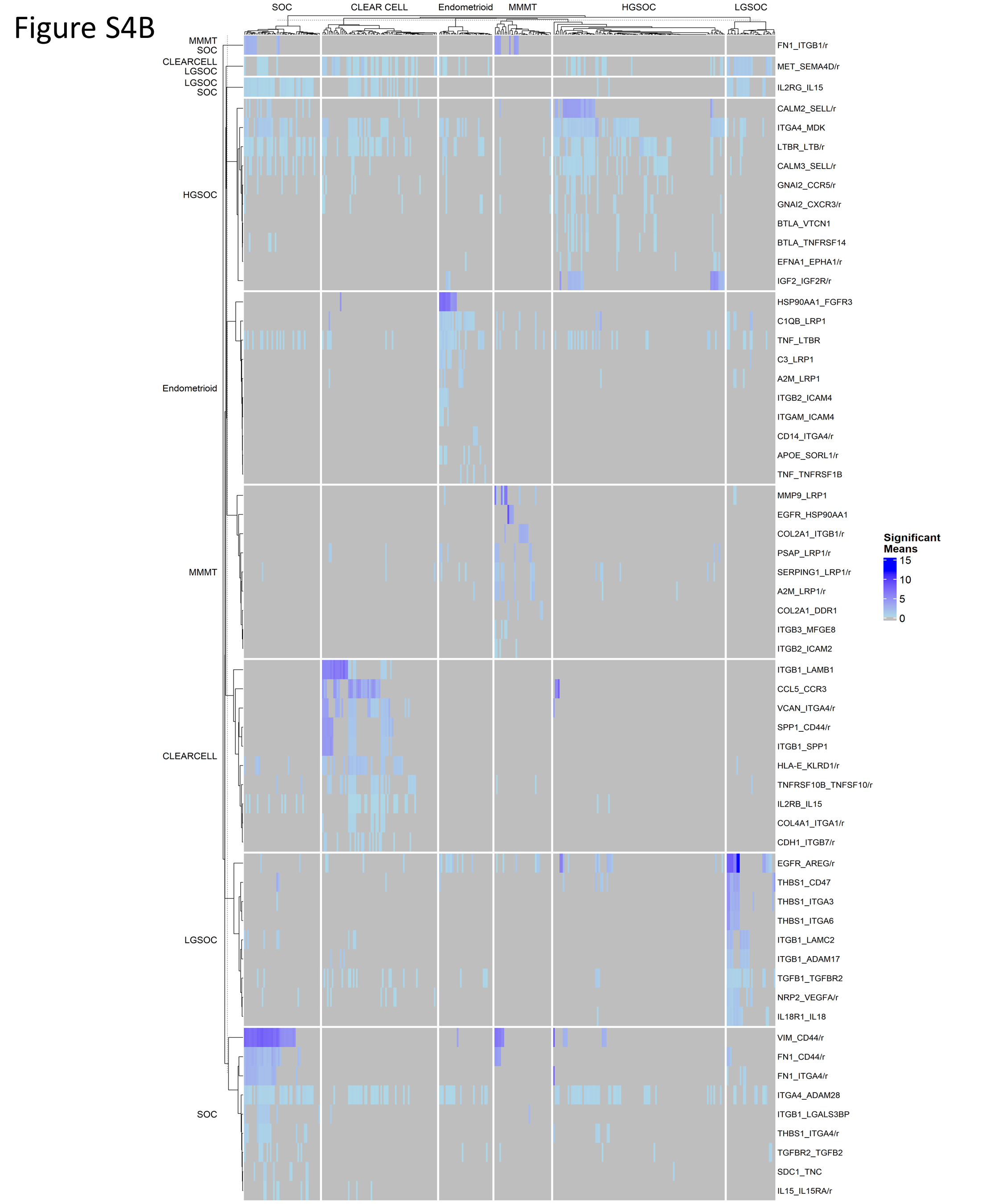

### Supplemental Figure 4C

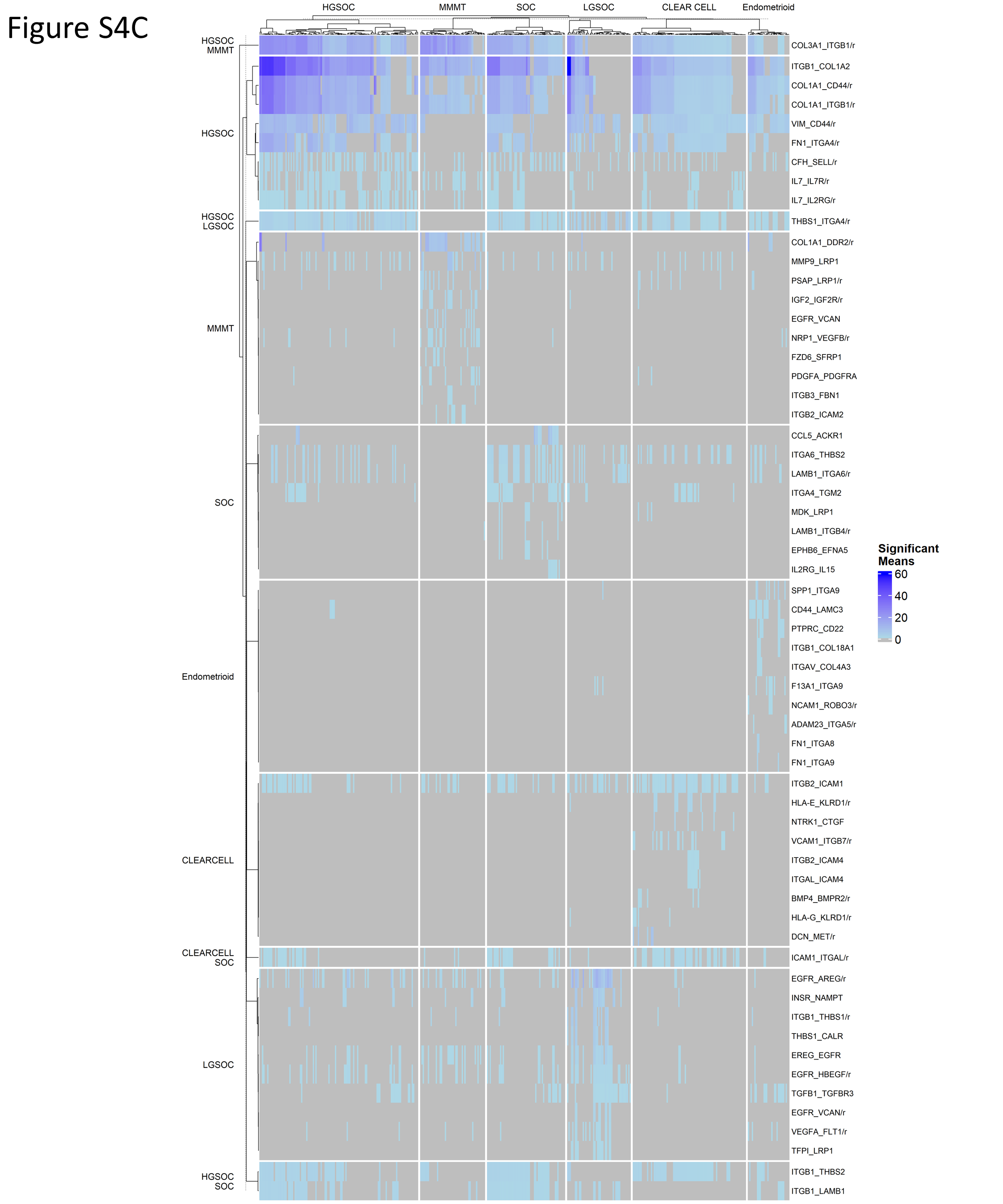

### Supplemental Figure 5

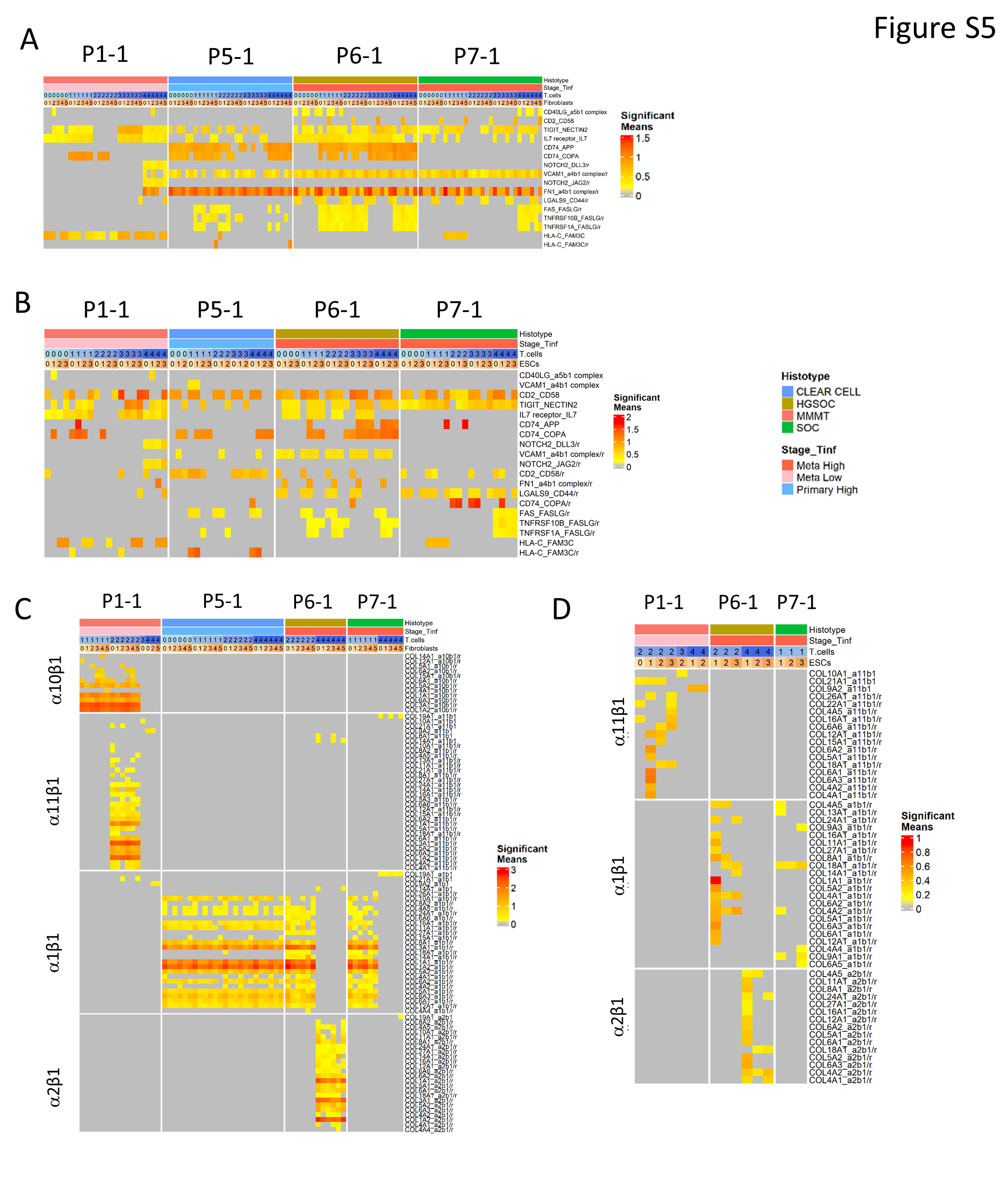
